# Scientific computing in the age of agentic AI: an exploratory field report

**DOI:** 10.64898/2026.07.29.741496

**Authors:** Jeremy Li, Alex Rubinsteyn, Sergey Feldman, Timothy O’Donnell, James M. Ferguson, Rob Patro, Ian Driver, Philip A. Ewels, Felix Krueger, Philipp Angerer, Ilan Gold, Jonathan Manning, Lukas Heumos, Mamad Ahangari, Varun Goyal, Hassan Masoudi, Brent Pedersen, Andrew Bai, Heng Li, Suyash Shringarpure, Gene Myers, Andrew Ho

## Abstract

Scientific computing has become a central component of modern scientific discovery. Yet many computational tools are developed by small, specialized teams under incentives that encourage the release of rapidly prototyped tooling without commensurate attention to engineering concerns, including performance and maintainability. These gaps are particularly visible in the life sciences, where the advent of high-throughput sequencing and molecular profiling has made the production and processing of datasets routine at scales that strain reliability and cost. Recently, LLM-based agents have become increasingly capable, with publicly available systems possessing both significant domain knowledge in many scientific fields and the ability to autonomously operate over complex and specialized codebases in pursuit of well-defined goals. Together, these developments create a practical opportunity for scientific computing. Many of the persistent weaknesses of the scientific computing ecosystem stem from technical debt and a shortage of sustained engineering labor and expertise. Here, we examine coding agents as a potential way to address these weaknesses: we present an exploratory field report of eight early case studies in the application of LLM agents to scientific computing across a range of project scopes, from lightweight maintenance tasks to full performance-oriented rewrites of scientific libraries, with a focus on the life sciences. Each of these case studies is accompanied by reflections from the individual or group responsible for the work, including lessons from the process. Overall, we find that the use of coding agents in scientific computing holds great promise for accelerating scientific research and increasing the reliability of critical systems, but that outstanding concerns remain, including responsibility and ownership for such projects, and we suggest collaboration and stewardship with existing maintainers when feasible.

## Introduction

Scientific computing is now a core pillar of modern research in both academia and industry.^1^ Across domains, increasingly large datasets are being generated, simulated, and collected; these data are then fed into computational workflows, with the goal of extracting quantitative evidence that supports scientific discovery and technological development. These trends can be seen in national cyberinfrastructure, where research allocations are measured in billions of core-hours, and in large scientific computing facilities that support thousands of researchers across universities, national laboratories, and industry.^2–3^ They are also visible outside national allocation programs in the increasing importance of computational science and research software engineering as recognized research roles across universities and industrial research organizations.^1,3–4^

Yet the libraries and tooling underpinning much of the scientific computing ecosystem are often produced under conditions that are quite different from those that support mature engineering practice in professional software organizations.^5–6^ Many tools are developed by small, specialized teams, especially in academic open-source settings, where software may be written to support a particular grant or method paper. The incentives governing funding, hiring, and career progression often reward new scientific results and publications more directly than their long-term maintenance.^4,6^ Indeed, in a recent review of the state of “research software” in the United States, Carver et al. describe such software as being typically developed by academics with “varying levels of training, ability, and access to expertise”, and identify recurring needs for engineering training and funding mechanisms which would encourage and credit software developers and maintainers.^4^ Industry and national laboratories also produce scientific software, but their development occurs under different institutional constraints; in practice, many open-source tools used in scientific practice originate in academic or laboratory settings where long-term maintenance is not always prioritized.^4–5^ This has resulted in a large amount of software that began as prototype implementations for particular papers becoming, in practice, long-lived shared infrastructure for a field, even though it was not originally developed with such in mind.^4–5,7^

These problems are particularly pronounced in the life sciences, where the introduction of high-throughput assays and large-scale data collection efforts have resulted in routine production of large datasets requiring complex and highly specialized computational pipelines at scales where reliability and performance become significant considerations.^8–9^ In genomics, this imbalance has been especially explicit: over the last 10-15 years, sequencing costs have fallen much faster than the cost of downstream analysis, resulting in the relative increase of compute-associated costs in the accounting of the total cost of a project.^8,10–11^ These compute-associated costs include not only the raw storage and compute-time costs but also the associated labor required to set up and maintain these highly specialized workflows.^9,11^

The resulting failure modes can extend beyond conventional software bugs; scientific code can be difficult to install, rely on deprecated dependencies, be weakly documented or under-tested, or be sensitive to undocumented parameters and workflow assumptions, thus impairing reproducibility and usability.^7,12^ In the best case, this results in wasted time on the part of users. In the worst cases, software, database, or workflow errors have challenged or invalidated published scientific conclusions, illustrating that implementation details can become methodological risks.^13–18^

Recently, LLM-based agents have begun to shift the application of AI models from being lightweight code suggestion tools toward goal-directed systems for repository-level software work. These systems combine LLMs, trained at scale on modern GPU infrastructure, with agent harnesses that enable them to autonomously pursue a well-defined goal, such as the implementation of a software tool. Recent models also show increasingly strong performance on expert-oriented scientific reasoning and data analysis benchmarks, suggesting that they may be useful in codebases where engineering tasks require specialized domain context. Repository-level benchmarks provide direct evidence for these capabilities: leading configurations on DeepSWE resolve a majority of 100-odd long-horizon tasks across five programming languages, while FrontierSWE extends evaluation to broader implementation and research-level problems, including in computational science and machine learning.^19,20^

This convergence creates a practical opportunity for scientific computing. Many weaknesses of the current ecosystem arise from the historical shortage of sufficient engineering labor and expertise in the development of scientific software, whereas such tasks are where coding agents excel. For example, coding agents can substantially reduce the effort of writing unit tests and high-quality documentation, tasks that are otherwise lengthy and tedious. They also speed up rapid prototyping of a method. Translating such prototypes into high-performance, production-ready code remains costly, however, as validation, security review, maintainability, and domain judgment still require substantial human effort. However, agents did not, for instance, fully complete any of FrontierSWE’s five from-scratch implementation tasks, suggesting that the reduction in engineering effort remains dependent on project scope and on whether the intended result can be clearly specified and validated.^20^

Here, we examine this opportunity through an exploratory field report of early case studies in which several independent groups applied LLM agents to scientific computing applications.

Across the eight case studies, we find that users applied coding agents across a range of project scopes, from relatively small, targeted patches intended to optimize or improve a limited surface area of a scientific tool to wholesale, performance-driven rewrites of algorithms in a different programming language.

We use these cases to identify recurring project types, validation burdens, and stewardship questions. We find that overall, users are broadly eager about the potential utility of coding agents, and find them to be a net positive to their work, while a persistent, outstanding question revolves around the best practices for communicating and collaborating with the original maintainers of the tool in question.

The projects fall into roughly six overlapping forms, listed below in approximate order of increasing scope:

- lightweight maintenance and packaging work,
- targeted optimization of existing functionality,
- compatibility migration between software frameworks while preserving released behavior,
- translation of tooling to new programming languages while preserving semantics,
- complete performance-oriented rewrites which involve major algorithmic changes,
- implementation of new tools, libraries, or new methodological capabilities within existing libraries.

Several projects span more than one category, and the degree of human oversight and intervention varied widely depending on the maturity of the existing implementation as well as the nature of the intended changes. We observe that the approach case study authors took to implementing their desired changes spanned a range of strategies, with certain groups taking the strategy of enforcing byte-level output equivalence, while others sought only to ensure logical equivalence.

Coding agents were most effective when the intended result could be checked against an external reference, such as byte-identical output, posterior agreement with an existing implementation, predictions from released statistical models, or existing test suites. Where exact equality was not expected, contributors instead used tolerances or domain-specific acceptance criteria fixed before evaluation. In some cases, divergence from reference implementations was found to result from inappropriate simplifications, altered logical flow, or errors made by agents during autonomous work. Authors also found that, while small synthetic workloads accelerated the pace of iteration, real-world data repeatedly exposed additional edge cases not present in simulated data.

In all but one project, human contributors concentrated their effort on the verification framework, often finding that they had to iteratively improve their framework as further work exposed additional failure modes. They concentrated their efforts on high-level control flow and system design, acceptance criteria, and the interpretation of discrepancies, serving more as product managers than as traditional software engineers.

## Case Studies

We present case studies submitted by eight groups who completed agentic coding projects in scientific computing, predominantly in computational biology, ranging from lightweight tasks such as modernizing the build process for an old but widely used genomic data I/O library to significant projects such as the wholesale liftover of a widely used but unmaintained RNA-seq aligner to Rust. The case studies were written by the contributing authors who executed the projects and are included in the **Appendix**.

For each case study, we requested that the contributor(s) describe the scientific context and motivating problem; the scope of the project; the role of the human in steering, testing, and validation; the evidence required to consider the result valid; the outcome; obstacles and failure modes; perspectives and lessons from the project; and links to the output artifacts. We standardized their organization and checked internal consistency and selected public artifacts where feasible, but did not independently reproduce every benchmark or validate every reported result. The authors of each case study are responsible for the accuracy and interpretation of its project-specific claims, including reported benchmarks and validation results. Unless otherwise stated, numerical results should therefore be interpreted as contributor-reported, case-specific outcomes rather than independently replicated estimates of agent performance.

Below, we summarize the findings, contents, and themes across the various case studies, focusing in particular on what contributors found agents to be able to perform autonomously and what they could trust them with, and what still required careful attention and a human in the loop.

**Figure 1:**
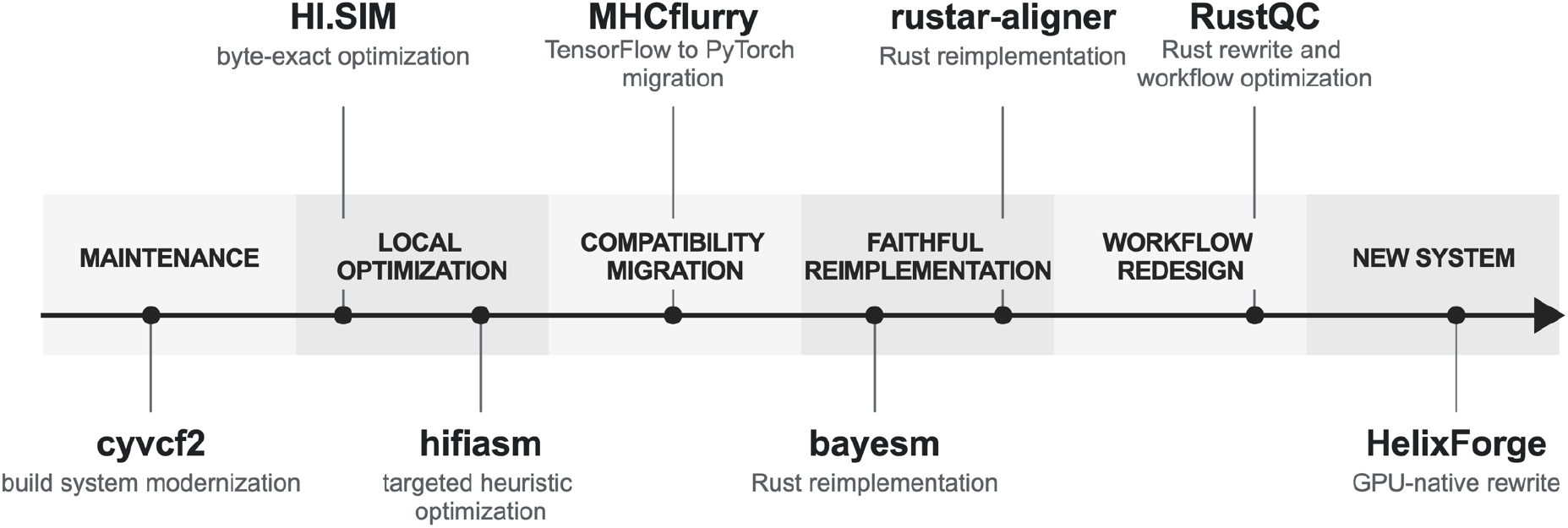
Approximate placement of the eight case studies by software surface affected and relationship to existing behavior, in approximate order of increasing complexity from left to right. For case studies spanning multiple projects, one representative project was chosen for illustrative purposes (rustar-aligner and RustQC).

The projects differed in their intended relationship to existing behavior and in the evidence available to evaluate their outcomes. Some aimed to preserve defined aspects of an existing implementation while improving performance or maintainability; others introduced new capabilities or substantially redesigned workflows, sometimes without an exact legacy reference. Several projects combined these aims, preserving selected outputs while altering implementation structure or introducing new workflow behavior. Accordingly, validation ranged from comparison with an existing implementation, where “equivalence” could mean byte-identical output or agreement on selected quantities, to independent evaluation against simulations, known answers, or prespecified acceptance criteria. Their numerical results are therefore reported as case-specific outcomes rather than direct comparisons of models or agent systems. Similar opportunities and validation problems nevertheless recur across the set.

Across the case studies, three properties were useful for organizing the projects: (1) the type of project; (2) the raw scope of the changes necessary, meaning the surface area affected; and (3) the validation target. The project types—for example, maintenance, targeted optimization, compatibility migration, reimplementation in a new programming language, workflow consolidation, and hardware-specific redesign—are meant to be descriptive and overlapping. The validation target depended on the project’s intended relationship to existing behavior.

Maintenance, targeted optimization, and compatibility migration generally sought to preserve defined scientific behavior, although the permitted implementation changes differed. For reimplementations, workflow consolidations, and hardware-specific rewrites, selected behavior was often preserved even as substantial portions of systems were changed.

Some case studies encompass multiple related projects. The Ewels contribution (Case Study C) centers on RustQC but also reports improvements to related quality control tools FastQC, FastQC-Rust, and Trim Galore; the Ferguson contribution (Case Study B) centers on rustar-aligner but also includes svb and kuva, new libraries implementing faster data compression and scientific plotting tools respectively. Table 1 summarizes the principal systems and selected related outcomes from each contribution, with full details provided in the Appendix.

**Table 1.**
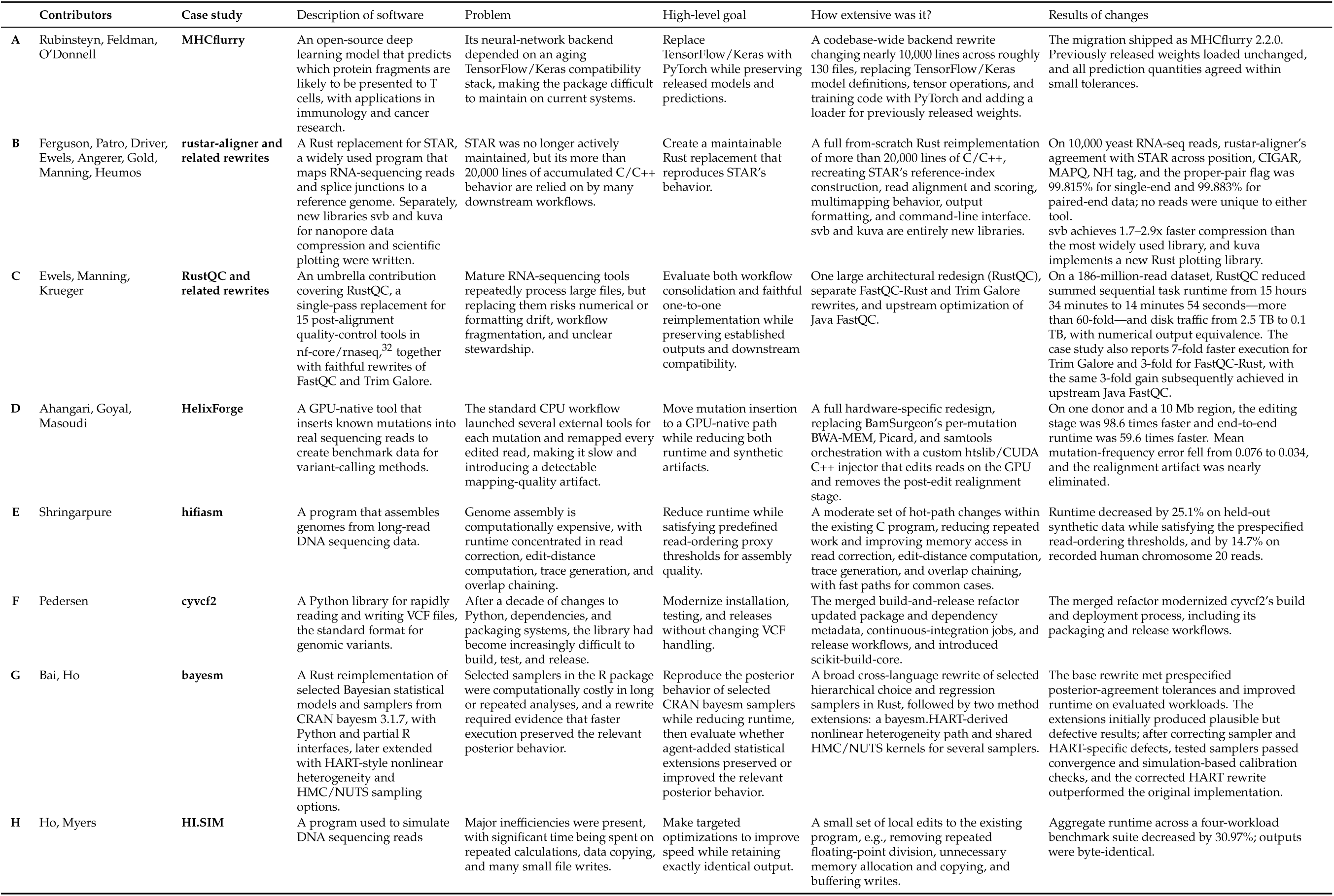
Summary of the eight agentic coding case studies. For each contribution, the table describes the software involved, motivating problem, goals, scope, and reported outcomes. Each case study can be found in full in the Appendix.

### Recurring Themes

We identified three recurring themes across all but one case study, reflecting the human’s primary role as a verifier and orchestrator. HI.SIM was the exception, with essentially no human intervention after the initial prompt. First, the burden of validation on the human in the loop increased with the software surface affected and with the extent to which a project introduced or altered scientific behavior. Smaller changes intended to preserve behavior were more straightforward to check through byte-level or exact numerical comparison, whereas broader or behavior-modifying rewrites that aimed to preserve an existing capability without requiring exact output equivalence required more complex comparisons across representative datasets, numerical results, and downstream workflows. Projects without an exact reference output required validation against simulated or synthetic data with known properties, or against other acceptance criteria fixed in advance.

Second, the projects generally proceeded through staged, feedback-driven iterations rather than as one-shot approaches. Contributors typically decomposed broad goals into iterated changes, and established intermediate benchmark/test harnesses against which agents had to test and revise changes. Initial implementations could often be produced rapidly, but resolving edge cases, subtle numerical differences, and failures that appeared only on realistic workloads frequently required substantially more iteration. Often, contributors found that completing the “last mile” of an implementation took the most work and effort.

Third, contributors reported that while agents could implement many well-scoped tasks, agent self-assessments did not provide reliable evidence of completion. In all but one case, contributors remained the principal adjudicators of success: they defined representative datasets and downstream quantities which served as required success gates, identified and investigated discrepancies across implementations, and determined whether the resulting evidence supported each intermediate and final claim. Across those seven cases, greater agent autonomy in implementation did not remove the need for human judgment; instead, it shifted human effort toward problem specification, validation design, and scientific interpretation.

## Discussion

In academic research, most software tools are written by small teams who primarily focus on making a clear methodological contribution; due to the cost and scarcity of professional engineering labor, factors such as adequate testing, code portability, and performance optimizations receive substantially less attention.^4–7^ Consequently, most academic code is difficult to run even immediately after release, let alone months or years following publication. The advent of coding agents, corresponding with dramatically lower software engineering costs, could make an optimized, tested-first release a routine expectation for academic code releases and allow for rapid modernization of established packages. Ideally, “out-of-the-box” releases should install predictably and expose consistent behavior across different system architectures.

Studies of code replicability indicate the scale of this opportunity. In a study of more than 9,000 published R scripts, 74% failed on the first run in a clean environment and 56% still failed after automated cleaning.^7^ Similarly, among 98 omics tools, 57.1% failed when the documented installation instructions were followed, 27.6% could not be installed after manual intervention, and an automatic failure required about 70 additional minutes of work on average.^12^ While these experiments are fairly limited, they provide a strong indication of the difficulty of basic code reuse. Considerable time and effort are expended on setting up the proper environment alone at the expense of performing scientifically interesting work.

For instance, consider 1,000 attempts to reuse research software with persistent failure rates between 27.6% and 56%. Under an illustrative scenario in which agent-assisted modernization prevented one quarter to one half of those failures, approximately 69–280 additional analyses would become runnable. Applying the reported 70-minute troubleshooting time would correspond to approximately 80–330 researcher-hours returned to analysis under these assumptions. At a fully loaded labor cost of $75–$150 per hour, this corresponds to $6,000–$49,000 across those 1,000 attempts, or $0.6–4.9M across 100 packages under the same assumptions. While these are very rough estimates, they illustrate the scale of researcher time that could be saved if modernization meaningfully reduced installation and execution failures. Additional benefits could arise from an enhanced ability to verify and extend published results, but we do not attempt to quantify these here.

Improved packaging, distribution, and reproducibility represent only one aspect of this opportunity. The case studies also suggest that agents can reduce the effort required to optimize existing software, making runtime and resource reductions more tractable, and can shorten the path from a scientific idea to a working implementation. By reducing the fixed cost of development, agentic coding may enable domain experts to prototype and implement tools that would otherwise have been substantially delayed or not developed at all. These benefits do not remove the need for validation and stewardship, but they expand the potential role of agents beyond maintenance and recovery of existing software.

### Validation and scientific correctness remain the bottleneck

Across all but one case study (HI.SIM), contributors converged on a broad validation strategy where the human in the loop defined and checked candidate agent implementations against a reference implementation or independently measurable target before considering something complete. In HI.SIM, the agent independently constructed the benchmark workloads and regression checks, with byte-identical output as the acceptance target. In the other cases, MHCflurry compared predictions from released weights within numerical tolerances, while more complex rewrites required more task-specific comparisons: hifiasm used a held-out workload and prespecified read-ordering thresholds; rustar-aligner compared read-level behavior against STAR; RustQC compared pipeline outputs; HelixForge evaluated mutation accuracy; and bayesm compared posterior population means under a prespecified tolerance. The bayesm extensions further illustrate that agreement on selected posterior summaries may be insufficient for new statistical functionality: first-pass HMC/NUTS and HART extensions produced plausible aggregate behavior, but additional convergence diagnostics, simulation-based calibration, and comparison against the original HART implementation exposed defects that required correction. The Ferguson auxiliary projects also illustrate settings in which automated checks were insufficient; for example, svb combined wire-compatibility tests with code review, while kuva paired numerical and structural tests with human inspection of rendered plots. Across all but one contribution (HI.SIM), developing and evaluating the validation framework was itself a substantial part of the human work.

Due to the inherent technical complexity of the optimization or rewrite projects, compilation and manual observation of ‘plausible’ outputs alone served only as very weak evidence of correctness. In practice, authors observed a number of subtle errors including changed numerical defaults, inappropriate memory allocation or streaming behavior, and silently skipped cases. Such failures can still result in plausible output that is subtly incorrect. For example, in rustar-aligner, progress beyond 90% parity required careful tracing of individual reads through both implementations to identify and resolve discrepancies. The validation harness itself can also be a source of errors; in HelixForge, an early false-positive strand-balance audit caused by downsampling led the agent to modify the GPU implementation even though the problem was in the auditing step. Careful human review is thus required at multiple levels of abstraction: the rewrite itself needs to be carefully checked, and the validation harness (which is often designed with the assistance of coding agents) itself also requires manual review.

The use of real-world data was found to be important for determining whether performance improvements transferred beyond small or synthetic workloads. For RustQC, validation across real public sequencing data showed that many edge cases surfaced only at realistic scale; minimal datasets were not sufficient. In hifiasm, the runtime reduction was smaller on recorded human reads than on the held-out synthetic benchmark, illustrating that performance gains can attenuate depending on the specific datasets. These cases suggest that agent-assisted optimizations should be evaluated on representative real-world workloads whenever feasible.

In all but one project, human judgment was invaluable in the continual evaluation and interpretation of quantitative improvements. While coding agents were capable of proposing and evaluating optimization hypotheses, the authors of those case studies had to repeatedly decide where to focus optimization pressure next and which higher-level strategy to attempt. This closely parallels the need for human-guided development of a validation strategy to ensure *technical* correctness; here, we also see that the application of human judgment is necessary to ensure the *conceptual* correctness of the end product.

### Stewardship remains a critical aspect

Faster and more reliable code is useful only if researchers are motivated to adopt it and if someone remains clearly responsible for ongoing maintenance. Commonly used scientific tools accumulate certain implicit conventions, such as the continual presence of undocumented (but important) behavior, understood only by their maintainers. Coding agents make it substantially cheaper to create a parallel implementation, but it often remains unclear whether that implementation should replace the original tool, be incorporated into it, or remain an experimental fork. In the absence of such clarity, there is a serious risk that the proliferation of inexpensive rewrites may simply divide users between superficially similar tools. Worse, the lack of standardization onto a single extension or rewrite may spread out valuable human oversight across an excess of similar projects; there is a serious risk that this diffusion of attention will result in ecosystems of software rewrites where no one rewrite is actually validated to an extent that permits real-world usage, even if concentration of efforts into a single project would have been able to produce a usable end product.

Authors of the case studies thus took different approaches to preemptively address the problem of code stewardship. The simplest approach is to incorporate patches directly into the original package. In cyvcf2, changes of varying scopes were prepared for the existing maintainers to evaluate, while the MHCflurry migration was completed and released entirely within the original project. The FastQC work described in the Ewels case study took a two-pronged approach—the tool was first rewritten in Rust and released as FastQC-Rust; following that, a number of performance improvements identified in that implementation were ported directly to the original, upstream Java implementation of FastQC. However, this is not always possible, such as with STAR; because the original program was no longer actively maintained, rustar-aligner was instead contributed to the scverse consortium,^33^ with pipeline integration and testing in nf-core,^32^ with the intention that future support is not dependent on a single contributor. These cases show that the appropriate outcome depends heavily on the state of the original project and the willingness of existing or new maintainers to assume responsibility for the new code.

Communication with existing maintainers should therefore begin well before a rewrite is ready for release. Contributors need to determine how to handle backwards compatibility, bug reports, licensing, attribution, and post-release maintenance. Rather than just being administrative details appended to the technical work, these questions are vital determinants of whether or not a prototype rewrite successfully evolves into a widely used codebase.

### Economic value of scientific software modernization

Agent-assisted modernization may also have economic value through reductions in compute and researcher time—in the following, we sketch a few stylized scenarios where we combine published workload parameters with explicit assumptions about adoption, runtime reduction, and unit cost to arrive at rough cost savings estimates. They should therefore be interpreted as illustrations of possible orders of magnitude rather than rigorous estimates of realized savings.

#### Direct runtime and compute savings

To illustrate the scale of potential runtime savings, we use a back-of-the-envelope scenario based on public RNA-sequencing records and the RustQC case study. The European Nucleotide Archive listed roughly 1.2M RNA-seq runs submitted in 2025, corresponding to ∼1M experiments and ∼1M distinct linked BioSample accessions.^21^ In the RustQC case study, RustQC processed a large paired-end human dataset of around 186 million reads through nf-core/rnaseq in 14 minutes 54 seconds, compared with 15 hours 34 minutes of sequential runtime for the original tools.^22^ Assuming all submitted RNA-seq runs undergo basic QC, replacing legacy tooling with RustQC would reduce the total number of QC-related compute-hours required for all annual ENA submissions by roughly 1.2–3.7 million CPU-hours.^23^ Actual realized savings would depend on workflow configuration and adoption, but the calculation suggests that runtime reductions in routine quality control could be substantial at current RNA-sequencing scale.

#### Ongoing software maintenance

Maintenance savings may be more plausibly estimated from the recurring work required to sustain a mature, widely used package. As yet another stylized example, NumPy, a foundational library in the scientific Python ecosystem,^24^ published 12 releases in 2025.^25^ During the same year, 326 merged pull requests had MAINT in their title, which we use as a narrow, title-based proxy for maintenance activity.^26^ Suppose that all of these changes were suitable for agent assistance and that, conservatively, such assistance saved two hours of human implementation time per change, without reducing the time required for review, testing, or release approval. This would save approximately 650 maintainer-hours per year, corresponding to roughly $49,000–$98,000 at a fully loaded labor cost of $75–$150 per hour. Given that pull requests vary widely in scope, and repository activity does not record the labor required to produce them, these numbers should be treated merely as illustrative. It nevertheless suggests substantial, recurrent value of agent assistance for a mature package, with the principal benefit being the redirection of scarce maintainer time toward compatibility decisions, code review, and stewardship.

#### Assisting and enabling new package development

Agentic coding also creates economic value when used to implement a new scientific software package from scratch. In the scenarios above, the counterfactual without agentic tools is a known workflow with measurable compute or troubleshooting costs. For an entirely new package, the relevant counterfactual may instead be a narrower prototype, a substantially delayed release, or no implementation at all due to limited software engineering bandwidth. The potential value here is therefore in unhobbling development in the frontier: reducing the fixed cost of implementation reduces the barrier to prototyping and experimentation, allowing domain experts to implement and disseminate tools which previously would have been time or resource prohibitive. The svb and kuva auxiliary projects, together with package-level systems such as RustQC and HelixForge, illustrate this channel in different forms.

At the same time, the case studies suggest that while agents can reduce the effort required to produce and improve implementations, they do not eliminate the costs of deciding what to build or remove the need for long-term responsibility for the package. The economic opportunity is therefore better understood as a reallocation of scarce expert effort from implementation toward specification, verification, and stewardship.

### Limitations

Several limitations should be noted. This exploratory field report is retrospective: the underlying projects were not commissioned for this study or conducted under a common protocol, and the case studies were collected from contributors after the work had already been undertaken. They therefore provide a narrow, selected cross-sectional view of current practice rather than a representative sample of agent-assisted scientific software projects or developers. Assessments of human effort, time saved, and economic advantage rely largely on contributors’ qualitative judgments rather than prospectively collected quantitative measurements. It is also worth noting that a number of other groups are pursuing more comprehensive programs of agent-assisted development, including but not limited to Fulcrum Genomics (high-performance genomics tooling, including Rust implementations of widely used sequencing QC tooling), Johan Henriksson’s group (workflow consolidation and semantically identical Rust rewrites of established scientific tools), and the Huang Laboratory (a full Rust rewrite of the Variant Effect Predictor and related Rust sequencing-quality-control tooling).^27–31^

## Conclusion

Across eight case studies, we describe how coding agents were used for a wide range of applications in scientific computing, ranging from maintenance and packaging to targeted optimization, language-level reimplementation, and hardware-specific redesign. Across the projects, coding agents provided engineering effort; in all but one, contributor effort focused on task definition, verification, and higher-level judgments. The cases and accompanying economic scenarios illustrate three plausible sources of value: reduced runtime and resource use, lower maintenance burden, and lower fixed costs for developing new tools. Realizing such value depends on careful validation as well as early coordination with maintainers where shared software is affected and the establishment of clear stewardship assignment for integration and maintenance. Overall, these case studies support the perspective that there exists great potential for coding agents in making scientific computing more durable, streamlined, and robust, but that the current bottleneck remains verification and validation.

## Author contributions

JL and AH conceived and coordinated the study and wrote the initial manuscript. AH, GM, MA, PA, AB, PE, FK, SF, JF, IG, VG, LH, JM, HM, TO, RP, BP, AR, and SS contributed the case reports. All authors reviewed and edited the manuscript.

### Acknowledgements

We thank Nils Homer, Tim Fennell, and Johan Henriksson for discussions about the topics considered in this article.

## Appendix: Individual case studies

### CASE STUDY A

### MHCflurry’s AI-Assisted Migration from TensorFlow to PyTorch

**Alex Rubinsteyn^1,2,3^, Sergey Feldman^4^, Timothy O’Donnell^5^**

^1^ Department of Genetics, University of North Carolina at Chapel Hill

^2^ Department of Computer Science, University of North Carolina at Chapel Hill

^3^ Department of Pharmacology, University of North Carolina at Chapel Hill

^4^ Allen Institute for Artificial Intelligence

^5^ Open Athena AI Foundation

### Summary

MHCflurry is a widely-used open-source immunology model which is useful for predicting T-cell targets in fields such as cancer immunotherapy, autoimmunity, and virology. MHCflurry’s neural network backend had drifted onto an unmaintained TensorFlow/Keras stack that resisted several attempts at migration. Between late January and mid-March 2026, two coding agents, Claude Code and Codex, alternated between contributor and reviewer roles to port the entire codebase to PyTorch. The agents changed nearly ten thousand lines of code across roughly a hundred and thirty files while preserving the ability to load previously trained weights unchanged. Outputs were validated against TensorFlow backend predictions on all predicted quantities. The result shipped as MHCflurry 2.2.0 and has been adopted smoothly by downstream users.

### Scientific context

MHCflurry^1,2^ is an immunology machine learning model that helps predict which parts of a protein inside a cell might become visible to patrolling T-cells outside that cell. Specifically, it predicts presentation of peptides on MHC class I molecules by considering both how a protein might be degraded into peptides and how well those peptides bind to diverse MHCs. Peptide-MHC complexes make the internal contents of a cell legible to the immune system, including viral proteins and tumor mutations. Predicting peptide-MHC presentation is thus a common step in research that has to prioritize possible T-cell targets out of a large pool of candidates.

We originally wrote MHCflurry in Jeff Hammerbacher’s lab at Mount Sinai because the standard predictors at the time were closed-source, slow, and couldn’t be retrained on additional data. Our motivating use-case was personalized cancer vaccine design: predicting which mutated peptides (neoantigens) actually get presented on tumor cells. MHCflurry has since become the most commonly used open-source MHC class I predictor, both by other groups working on cancer immunotherapy^3^ and in other domains of immunology such as virology.^4^

### Problem or opportunity

MHCflurry development began in 2015 using the Keras neural network library, when its default GPU backend was Theano.^5^ In the early 2010s, Theano had been a revolutionary tool for deep learning research in Python but had notoriously messy internals which generated many difficult-to-debug errors and error messages.

We eagerly switched the backend in MHCflurry’s use of Keras to TensorFlow^6^ in 2017, a few months before Theano’s own developers announced they were no longer maintaining the project. TensorFlow, unfortunately, proved to be an unstable substrate, and its frequent breaking changes imposed a maintenance burden on our project.

TensorFlow 2.0 (2019) then changed its default execution model. MHCflurry couldn’t easily move to the new eager-execution mode, so we stayed pinned to TensorFlow 1, annoying some of our users. We eventually upgraded to TensorFlow 2 in 2020, but only by dropping into a compatibility mode. For fear of future breaking changes, we didn’t try to keep up with newer releases of the dependency, further annoying some users.

As deep learning matured as a field, the center of gravity of the open source models moved to PyTorch.^7^ We wanted to drop both TensorFlow and Keras entirely and rebuild the models in PyTorch, ideally while keeping the ability to load previously trained weights without significant changes in predictor outputs. Since we were already going to be performing intrusive surgery on the numerical dependencies, a secondary goal was to also modernize the rest of the numerical stack (Pandas and NumPy).

### Agentic intervention

The intervention consisted of a backend rewrite/migration of a mature, numerically sensitive scientific library, followed by targeted feature development (multi-GPU training) on a pre-release branch.

### Human role

The human contribution was primarily steering and orchestration:

- Specification of acceptance criteria and creation of a small validation dataset.
- Deciding to add a second agent (Codex) as reviewer/contributor once subtle numerical problems persisted after prolonged work with just Claude Code.
- Choosing the pattern of agentic work: develop with one harness, review with the other, and switch roles when work stalled or moved unproductively.

### Validation and evidence

PyTorch outputs were checked against the TensorFlow backend predictions across 315 allele-and-peptide combinations and agreed to a very small error tolerance across all predicted quantities (affinity, processing, presentation, and their percentile ranks). A loader was added so the new code reads the old trained weights unchanged, so equivalence could be tested on the released models rather than on freshly retrained ones. Post-release, the upgrade has been adopted smoothly by downstream users, consistent with the published claim that 2.2.0 loads the same weights and produces equivalent predictions with no workflow changes.

### Outcome

The migration shipped as MHCflurry 2.2.0, the first stable release on a PyTorch backend, and has been adopted without friction downstream. Development has continued in the same two-agent style on a pre-release 2.3.0 that modernizes the pan-allele training pipeline with device-resident, multi-GPU training which cuts the most intensive training step (fitting a large ensemble of affinity predictors) from about a week to about an hour on an 8xA100 node.

### Obstacles and failure modes

The recurring blocker across every attempt, human and agentic alike, was divergent behavior between networks that looked superficially identical. Small discrepancies originated from different default parameters (e.g. padding/edge modes used for convolutions), genuine numerical differences between the TensorFlow (or NumPy) implementations of an operation and its PyTorch equivalent, or from subtle mistakes while porting neural network specifications.

These regressions were numerous and subtle enough to deter previous port attempts by humans and to slow an agentic porting effort in early 2025.

That early attempt is worth describing as good context for the eventually successful later effort. One of us (S.F.) first tried to migrate MHCflurry with aider,^8^ an early open-source CLI-centric agent harness, using Claude 3.5 Sonnet and OpenAI’s o1 model. Though this effort did successfully port one component of MHCflurry over a week of work, it was then abandoned because of the perceived immaturity of the agentic coding process. When closing his pull request, S.F. wrote:

> *Why 200 commits? Because I did this almost entirely with aider… I have learned a lot about how incredibly naive AI-code generation is if you just let it tell its own stories without constant questioning, re-questioning and demands for more tests/debugging/logging/analysis.*

With a year’s hindsight, S.F. places the bottleneck on model capability rather than tooling:

> *I think this was practically impossible until Opus 4.5 (or its equivalent GPT model). The sophistication of the available harness was likely not the primary bottleneck. The error compounding was too severe, and the model’s intelligence/capacity was too low to complete this task at the level of autonomy I hoped for.*

Even in the later efforts using better harnesses and models, we found that agents excel at mechanically migrating APIs across dozens of files but still litter generated code with subtle errors. Agentic code reviews are not always sufficient to reach convergence but might instead nudge the agent towards unnecessary expansion of the code surface, which in turn introduces even more subtle bugs. It took a great deal of questioning and sanity checking by a human to slowly achieve near-equivalence of predicted outputs.

### Lessons and reflections

- Adversarial pairing beats a single agent. Letting Claude Code and Codex alternate between contributor/reviewer roles seemed to escape plateaus since the two agents caught different errors in their reviews.
- Define acceptance criteria. A concrete acceptance criterion gives the agentic loop a target for more autonomous iteration. In our case we required small relative and absolute differences across all predicted quantities for a predefined allele/peptide dataset.
- Preserve backward compatibility. Loading publicly released model weights unchanged made equivalence testing against a ground truth easier.
- Agentic coding is great for maintenance. The ability to do this kind of unglamorous, labor-intensive upkeep is often what keeps an open-source scientific project alive rather than decaying.

### Artifacts

|  |  |
| --- | --- |
| Source repository | openvax/mhcflurry |
| Documentation | MHCflurry documentation |
| Merged pull request | PR #260: "TF to PyTorch (an attempt by robots where humans have failed)" |
| Prior attempt | Sergey Feldman's abandoned aider PR #249, "Torch support part 1" |
| Release | MHCflurry v2.2.0 (first stable PyTorch-backend release) |
| Prerelease | MHCflurry v2.3.0rc6 (device-resident / multi-GPU training-pipeline work) |

### References

1. O’Donnell TJ, Rubinsteyn A, Bonsack M, Riemer AB, Laserson U, Hammerbacher J. MHCflurry: open-source class I MHC binding affinity prediction. *Cell Systems.* 2018;7(1):129–132.e4.

2. O’Donnell TJ, Rubinsteyn A, Laserson U. MHCflurry 2.0: improved pan-allele prediction of MHC class I-presented peptides by incorporating antigen processing. *Cell Systems.* 2020;11(1):42–48.e7.

3. Hundal J, Kiwala S, McMichael J, et al. pVACtools: a computational toolkit to identify and visualize cancer neoantigens. *Cancer Immunology Research.* 2020;8(3):409–420.

4. Leddy OK, White FM, Bryson BD. Leveraging immunopeptidomics to study and combat infectious disease. *mSystems.* 2021;6(4):e00310-21.

5. Theano Development Team. Theano: a Python framework for fast computation of mathematical expressions. Preprint. 2016.

6. Abadi M, Agarwal A, Barham P, et al. TensorFlow: large-scale machine learning on heterogeneous distributed systems. 2016.

7. Paszke A, Gross S, Massa F, et al. PyTorch: an imperative style, high-performance deep learning library. *Advances in Neural Information Processing Systems 32.* 2019. p. 8024–8035.

8. Aider. Aider: AI pair programming in your terminal. Accessed 2026-07-14.

### CASE STUDY B

### Faithful rewrites, foundational primitives and the limits of agent self-verification: three Rust projects with coding agents

**James M. Ferguson^1^, Rob Patro^2^, Ian Driver^7^, Phil Ewels^3^, Philipp Angerer^4,6^, Ilan Gold^4,6^, Jonathan Manning^3^, Lukas Heumos^5,6^**

^1^ Garvan Institute of Medical Research

^2^ University of Maryland

^3^ Seqera

^4^ Institute of Computational Biology, Helmholtz Munich

^5^ NVIDIA Corporation

^6^ scverse

^7^ Independent researcher

James Ferguson contributed to all three projects; the remaining contributors are specific to rustar-aligner. Claude Code (Sonnet 4.5/4.6) was used throughout.

### Summary

This case study explores the role of coding agents in the creation of Rust tools and libraries across a range of interventional scopes. Three tools were either built or rewritten with the aid of coding agents. The consistent finding is that while agents reduce the cost of engineering labour, their role remains limited by the need for human guidance and judgement, reflecting a need for partnership over delegation, in order to deliver a comprehensive project. The type and breadth of human verification differed by project and are characterised in this case study. The three projects include a faithful rewrite of a large unmaintained aligner (rustar-aligner); a foundational SIMD codec primitive with a novel algorithmic extension (svb); and a scientific plotting library/CLI with over 60 plot types (kuva).

#### rustar-aligner

**Project contributors:** James Ferguson (Garvan Institute of Medical Research, Rust rewrite), Rob Patro (University of Maryland, project lead/caps-sa index), Ian Driver (Independent researcher, rustar-solo), Phil Ewels (Seqera, nf-core), scverse/nf-core community members: Philipp Angerer (Institute of Computational Biology, Helmholtz Munich), Jonathan Manning (Seqera), Ilan Gold (Institute of Computational Biology, Helmholtz Munich, AnnData integration), Lukas Heumos (NVIDIA Corporation). Original tool: STAR, by Alexander Dobin, who has been approached to participate in an advisory capacity. Maintained by scverse with pipeline integration and testing in nf-core.

STAR is a highly used RNA-seq aligner whose active development and active maintenance seem to have ceased. Given the continued widespread usage of STAR, contributors from two communities, scverse and nf-core, want to ensure that STAR is sustainably maintained and developed. To make contributions easier while improving memory safety, speed, and efficiency, we set out to rewrite STAR in Rust with the help of coding agents. The resulting software, termed rustar-aligner, is a drop-in replacement: the same arguments, and close to identical output. It reaches 99.815% single-end and 99.883% paired-end tie-adjusted parity with STAR, with no reads mapped by one tool and not the other, and a suffix array byte-for-byte identical to STAR’s. Closely reproducing the results of a reference implementation is a useful first step in winning a user community’s confidence. Recent algorithmic advances then open clear opportunities for improvement, which later development will pursue.

#### svb

**Project contributors:** James Ferguson (Garvan Institute of Medical Research)

svb is a pure Rust StreamVByte library: integer compression with delta and zigzag transformations across 16-, 32-, and 64-bit widths and SIMD backends for AVX2, SSSE3, and NEON. It is wire-compatible with existing StreamVByte data and runs consistently 1.7 to 2.9x faster than streamvbyte64, the most used existing crate. I wrote svb for use by native Rust nanopore signal file libraries (it is used in my pod5lib and slow5lib crates and was built to be wire-compatible with existing pod5/slow5 files). Beyond the reimplementation, the work produced a novel algorithmic extension: a fused, multi-stream decode method called VBZ2/VBZK that breaks the serial carry-chain limit normally treated as fixed, opening a path to near-linear multicore decompression at a tiny compression-neutral format cost.

#### kuva

**Project contributors:** James Ferguson (Garvan Institute of Medical Research) kuva is a new scientific plotting library in Rust. The first 11 plot types and library architecture were written by hand. Afterwards, agents were used to accelerate development and completeness of the project. It has now grown to over 60 plot types, each shipped with documentation, examples, tests, and a CLI tool with terminal plotting. It has SVG, PNG, and PDF backends, and minimal dependencies for a fast and lightweight library. It has gained organic adoption: independent conda-forge and AUR packaging, various crate dependents, and multiple contributions from users.

### Scientific context

#### rustar-aligner

STAR remains the alignment tool of choice in a wide body of RNA-seq analysis workflows.

A reimplementation must build confidence by replicating STAR’s results as closely as possible. “Close enough” is not acceptable for a tool at the centre of mission-critical production pipelines.

The original is over 20,000 lines of C/C++.

#### svb

StreamVByte is the integer compression method used before going through zstd in nanopore signal file formats (pod5, slow5). Every tool that reads raw nanopore signal has to do the decompression of the signal using the available libraries, at the scale of millions of reads per sequencing run. It is the algorithmic bottleneck in decompression, and most implementations reach the theoretical maximum throughput with SIMD. A primitive codec like svb doesn’t receive much attention or engineering effort once this theoretical maximum has been reached (agents were sure this couldn’t be beaten), particularly given the brutal effort-to-reward ratio of handwritten SIMD code.

#### kuva

Bioinformatics needs publication-quality, scriptable plotting that runs in pipelines without a Python/matplotlib or R/ggplot2 dependency stack. kuva covers many genomics-specific plot types (Manhattan, UpSet, phylogenetic trees, synteny diagrams, Kaplan-Meier curves, ROC/PR curves, clustermaps) alongside standard plot types. There was also a need for a better Rust library that made plotting easier, with an intuitive API.

### Problem or opportunity

#### rustar-aligner

Despite STAR no longer being in active development, its continued widespread adoption made ongoing support important for both the scverse and nf-core communities. To ease contributions, a more modern implementation in Rust was proposed, though, given the size and complexity of the original, a manual rewrite was not a practical consideration. However, the availability of coding agents capable of compressing that timeline and effort considerably presented an opportunity to generate this new implementation and provide a good basis for ongoing maintenance and development.

#### svb

A fast, wire-compatible StreamVByte codec in both 16- and 32-bit widths was needed for use in VBZ pipelines for pod5lib and slow5lib Rust rewrites. During the development of svb and the VBZ methods (delta, zigzag, svb encoding/decoding), the main bottleneck was the delta serial prefix sum carry chain. However, this bottleneck was an algorithmic one, not an implementation one. The opportunity was a library that contained a more complete set of svb methods, including the delta and zigzag variants, and to find a way to get past the delta speed limit.

#### kuva

Plotting is where bioinformatic tooling tends to be weakest in Rust: The scientific plot vocabulary isn’t well covered, and the usual problem with single-maintainer projects is that features outrun the documentation and tests, and the project becomes unmaintainable long term. The opportunity was to expand the breadth of a plotting library without that decay, keeping docs, CLI, and test coverage moving in step with each new plot type.

### Agentic intervention

#### rustar-aligner

Human-steered agent for a fully faithful reimplementation (port by behaviour) of an existing tool whose maintenance has lapsed, written from scratch rather than transpiled.

#### svb

Human-steered agent for a new implementation of a foundational primitive, with rapid SIMD iteration, along with exploratory algorithm design that produced a novel extension.

#### kuva

Human-directed agent to implement new features at scale, expanding an existing hand-written core to many new plot types. The hand-written core provided a scaffold for the agent to use for new plots, making the API surface coherent across plot types.

### Human role

#### rustar-aligner

A rewrite of this kind cannot be delegated to an agent with a single instruction such as “rewrite this in Rust, no mistakes.” The two languages differ in ways that demand deliberate decisions on architecture and library selection to keep the resulting code maintainable. For this reason, the port was written from scratch with agents rather than through automated tooling such as c2rust. Architectural and library decisions, along with the testing strategy and methodology, were determined manually, while the agent handled implementation and iteration. The decisive contribution was the verification methodology. A further requirement was keeping the agent progressing through regressions rather than reverting prior work, which proved a persistent challenge but was essential to achieving parity. These rules were encoded directly into the agent’s standing instructions, and the agent was guided toward parity throughout.

#### svb

Agents are well suited to SIMD iteration, normally the most difficult aspect of building a library such as svb. Writing SIMD by hand is slow and error-prone, and exploring multiple approaches to identify the optimal one can be time-consuming. With the agent, a full iteration and testing loop could be completed in roughly an hour. Translating the final solution from AVX2 to SSSE3 and NEON was then straightforward for the agent, although writing and testing such translations is typically difficult.

This iteration speed made it inexpensive to try different approaches, and led to the observation that the zigzag and StreamVByte steps could be folded into the gaps of the delta carry chain, rendering them nearly free and yielding a substantial speedup without loss of compatibility. Many other approaches were attempted that performed poorly, but the speed of iteration with an agent kept the trial-and-error phase relatively cheap. The same efficiency made it possible to sketch the VBZK multi-stream method, which achieves a near-linear speedup with additional threads.

#### kuva

Each plot was scoped individually, along with the API conventions and per-plot semantics, and every rendered plot was reviewed visually. Each test produces a plot that can be inspected and used to guide the agent toward correct output.

### Validation and evidence

#### rustar-aligner

STAR itself is the ground truth. The most important step is a testing framework with appropriate data for checking and classifying differences between STAR and rustar-aligner. Parity was measured on position, CIGAR, MAPQ, NH tag and proper-pair flag against STAR 2.7.11b on an identical index and arguments, using 10k yeast RNA-seq reads. Parity testing yielded 99.815% SE and 99.883% PE tie-adjusted parity, 0 STAR-only and 0 rustar-only reads, 0 MAPQ inflations or deflations, 0 NH differences, and a suffix array with 10,862 entries that is byte-for-byte identical to STAR’s. rustar-aligner has 396 passing tests. Some output is legitimately random, for example the choice of which equal-scoring multimapper is marked primary, which is random and suffix-array-order dependent. Filtering those out defines the real accuracy ceiling.

Additional validation was carried out through trials of rustar-aligner within the nf-core/rnaseq workflow, exploiting the in-built test framework there. That surfaced additional differences that impacted downstream tooling. Examples included mis-bucketed unmapped-read counters, a +33 offset on the BAM QUAL field that doubled the reported average quality, and dropped transcriptome mate fields that degraded expression estimates generated by Salmon using rustar-aligner alignments.

#### svb

Ground truth is wire compatibility: svb’s output is byte-compatible with existing StreamVByte data and checked against fixed external formats (pod5 and slow5 files). Performance was measured in a head-to-head benchmark against streamvbyte64 v0.2.0 on identical inputs (AVX2, GitHub Actions CI, throughput in GB/s of input integers):

- U32Classic decode: 2.34x to 2.88x faster across 128/1024/8192-element slices (14.1 vs 4.89 GB/s at 8192).
- U32 encode: 2.68x to 2.85x. U64Coder1248: 1.69x to 2.04x (smaller gap, as 8-byte elements expose less per-control-byte parallelism).
- Fused VBZ decode: 3.68 GB/s, 1.52x the 3-pass pipeline (svb->zigzag->delta), reaching 99% of the delta-alone limit.
- VBZ2 2-chain decode: 5.62 GB/s, 1.53x over single-chain fused and 2.32x over 3-pass pipeline (svb->zigzag->delta).

Wire compatibility is verified in CI testing using round-trip encoding and decoding between svb and streamvbyte64 for the various codecs.

#### kuva

There is no external ground truth for a plot the way the other 2 projects have. A plot is correct when it is both numerically right and readable, where readability is a visual judgement. Validation comes in two parts. Unit tests check the numerical and structural parts of a plot, and every test renders a plot, with additional smoke tests and terminal test plots, which are checked by eye.

### Outcome

#### rustar-aligner

The principal outcome is a behaviourally faithful, memory-safe and memory-efficient Rust implementation of STAR (rustar-aligner), developed and maintained by scverse with pipeline integration and testing by nf-core. rustar-aligner is MIT-licensed to match the original. With near-parity reached, extension has begun. We replaced the suffix-array builder with caps-sa to cut genome-index build time by a considerable fraction and to make build memory proportional to the input text size rather than the final output size (reducing build memory by a factor of 3 on a dual-strand human genome index, for example), while maintaining identical byte-for-byte indexes. We are also making arguments more user-friendly, for example by having --limitBAMsortRAM accept both raw bytes and human-readable values like “64G”. The adoption of rustar-aligner by scverse, an established community with a substantial Rust contingent, is another important outcome. This will allow rustar-aligner to have institutional continuity rather than depending on a single maintainer. Multiple contributors have joined as maintainers since reaching near-parity, supporting this objective.

#### svb

A wire-compatible codec ∼1.7x-2.9x faster than the most used library (streamvbyte64), a Rust VBZ encode/decode library for nanopore signal file handling, plus a novel parallel-decode extension with a path to multi-core signal decompression at a compression-neutral cost. Full benchmarks, documentation, and a polished crate across three SIMD targets.

#### kuva

A library and CLI with more than 60 plot types, complete docs, and multiple backends, built in a fraction of the time a solo developer would need to get to the same finished state, where docs/packaging/tests would usually get deferred. Since release, multiple users have submitted issues and pull requests, adding features and their favourite plot types. Some did so using various agents; others did so by hand.

### Obstacles and failure modes

#### rustar-aligner

This project produced the clearest examples of agent limitations in my work.

At around 90% parity, the agents stalled. The remaining differences consisted of several layers of bugs stacked upon one another, so that any single change caused a regression in testing; the agent would then revert its work, unable to progress. The breakthrough came from allowing the agent to modify the STAR source alongside rustar-aligner, adding debug output to trace individual reads through both alignment pipelines. This pinpointed where the two diverged, and the layered bugs could be identified and resolved one at a time. Permitting regressions while following this path—trusting the overall process rather than the intermediate tests—was essential to reaching 99.8+% parity.

Other recurring failure modes were observed. One was the use of >= where STAR uses >: a subtle error that did not always trigger and was nearly invisible in read outputs, exemplifying the case in which code that compiles and appears correct is not. The agent also repeatedly tripped on long-read-only #ifdef blocks in STAR’s C++, making algorithm decisions based on code paths not active for short reads; the source-tracing method largely resolved this, since edits to dead code produced no change and forced the agent to investigate further. More generally, agents tend to decompose a complex, tightly coupled algorithm into atomic pieces and then cannot recover the accuracy that the monolithic design achieves. Some complexity is irreducible, and the agent fights it.

#### svb

During development, the agent repeatedly attempted to implement simplified methods that were not correct, requiring continual steering back toward completing the work as specified rather than approximating it. These attempts were caught by reading the code at each iteration rather than through the test frameworks, as the agent wrote tests that “passed” and appeared plausible but were not valid. The novel components, VBZ2/VBZK and Fused VBZ logic itself, were handled well once specified; the difficulty lay mostly in standard codec correctness. The agent’s strengths and weaknesses here are two sides of the same coin: it iterates on SIMD quickly enough to be transformative, yet cannot, on its own, distinguish a fast, correct method from a fast method that merely appears correct.

#### kuva

kuva demonstrates the clearest failure mode: agents simply cannot self-verify yet. The agent will assert that a plot looks correct, but on visual inspection it is severely flawed: overlapping labels, an incorrect axis, an element flipped or rotated. It has no perception of the small details that make a plot readable. This failure is so common and so complete that the only reliable way to handle it is structural. Every test in the cargo suite renders a plot, over 900 so far, and each is examined visually for regressions or omissions across all plot types.

### Lessons and reflections

Across all three projects, the same shape recurs: coding agents reduce the cost of engineering labour, but the work only succeeds through partnership over delegation. The agent serves as an implementer and fast iterator, with the human holding the architecture, judgement, and verification. Whenever work was delegated to the agent, it produced plausible but incorrect output; when the agent was used in partnership—steered continuously and backed by human judgement about correctness—it produced tools we are willing to take responsibility for.

The verification that stays human differs by project. For rustar-aligner it was identifying the need to trace single reads through two pipelines to separate layered bugs. For svb it was reading the code on every iteration because the agent could not be trusted to implement correct methods. For kuva it is the visual question of whether a plot is readable or not, which no model can currently judge effectively, leading to over 900 plots being manually checked by eye before a release. The common thread is that agents are strong at execution and iterating fast against a concrete target, and weak wherever correctness is not well defined: where it is silent and only shows up downstream (rustar-aligner), where the agent’s own tests certify the wrong answer (svb), or where the judgement is visual (kuva).

Building svb highlighted where human insight lives during the development cycle. The agent was certain the SIMD throughput limit had been reached. Getting past that point required iterating through a set of trial-and-error methods, eventually leading to the discovery that svb+zigzag takes roughly as many cycles as delta, and so you can get all three stages in one pass through memory. The agent made testing this idea cheap to implement and measure. Fast iteration lets us quickly build up to the limit and explore the edges.

The case studies here concern rewrites, primitives, and feature breadth, but the same capabilities point to a broader opportunity. The svb work showed agents iterating on hardware-specific vector code fast enough to be transformative, translating a tuned solution across three SIMD targets that are normally slow and error-prone to write by hand. Much of the performance-critical work in scientific computing has this character: the barrier to substantial speedups is often the specialized expertise required to target the hardware. Agents may lower the cost of accelerated computing in the same way, making it more tractable to adapt performance-critical code to architectures such as GPUs, bringing speedups within reach for domain scientists who would not otherwise take on that work. As with the rest of this study, the constraint would remain human: the same verification burden applies, since a fast kernel that is silently wrong is exactly the failure mode svb surfaced.

Finally, agents lower the cost of rewrites and reimplementations to the point where the constraint is no longer effort but judgement and provenance. How a tool is made matters far less than whether someone takes responsibility and maintains it. rustar-aligner found a home in scverse before the first near-parity version was created; kuva, largely built with agents, is healthy because of user participation through issues and pull requests, some also written with agents. Our rule of thumb covering both rewrites and new tools still applies: don’t build it if you aren’t prepared to support and maintain it, agents or not.

### Artifacts

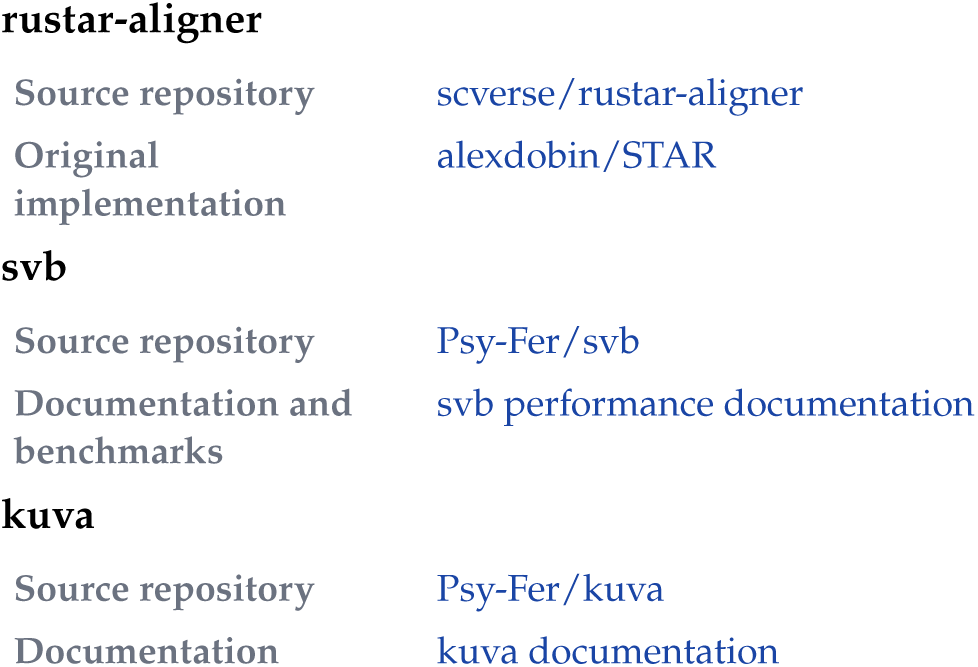

rustar-aligner
|  |  |
| --- | --- |
| Source repository | scverse/rustar-aligner |
| Original implementation | alexdobin/STAR |
| svb |  |
| Source repository | Psy-Fer/svb |
| Documentation and benchmarks | svb performance documentation |
| kuva |  |
| Source repository | Psy-Fer/kuva |
| Documentation | kuva documentation |

### CASE STUDY C

### Agent-assisted rewrites of RNA-sequencing QC tools: RustQC, FastQC-Rust, and more

**Philip Ewels^1^, Jonathan Manning^1^, Felix Krueger^2^**

^1^ Seqera

^2^ Altos Labs

### Summary

Nextflow is the leading workflow manager in bioinformatics; nf-core/rnaseq,^1^ the most popular pipeline in its nf-core community, is run by thousands of groups and companies worldwide to process RNA sequencing data. I rewrote its QC tools into a single binary called RustQC, achieving a 60x speed-up that reduces QC processing time for one sample from 15 hours to 15 minutes. My experience of this process and the agentic tool rewrites that followed led me to create rewrites.bio, a set of principles for AI-assisted rewrites. Agentic coding makes the technical debt we face in bioinformatics a tractable problem, but we as a community need to be cautious of fragmentation, trust and verification.

### Scientific context

Nextflow is an open source workflow manager for chaining specialised tools into reproducible, portable pipelines, with a language and execution layer that keep results consistent across systems. The nf-core community has grown alongside it, with 14,000+ scientists collaborating on a curated set of pipelines and workflow components.

Quality control is a critical step in almost every sequencing analysis, and for RNA-seq the community standard is the QC subworkflow in nf-core/rnaseq. That subworkflow runs fifteen separate tools: dupRadar, featureCounts, eight RSeQC modules, preseq, three samtools commands, and Qualimap. Most have been part of nf-core/rnaseq since I first wrote the workflow in 2016. They are well established and provide outputs now built into the results-verification systems of hundreds of organisations. Faster tools exist in some cases, but the originals carry momentum from their time in the community, making them costly to switch away from. As such they have remained mostly unchanged for over 10 years.

Other key parts of the pipeline are raw sequence QC (FastQC), quality- and adapter-trimming (Trim Galore) and alignment to a reference genome (STAR). Together they take raw sequencing data and produce read count matrices suitable for differential gene expression analysis and more.

### Problem or opportunity

dupRadar was one of the first QC tools added to nf-core/rnaseq. It is an R library written by Sergi Sayols Puig during his PhD that measures the complexity of RNA-seq libraries, warning of insufficient input quantity or quality. RNA-sequencing was mostly small-scale by today’s standards at the time, and computational efficiency was not the highest priority. Under the hood, dupRadar runs the Subread FeatureCounts tool four times per sample, with and without duplicates and multi-mapping reads. A similar story extends across the QC tools: each iterates over the input data and writes its own intermediate files before producing summary statistics. Taken together, for a single 10GB BAM file they generate nearly 2.5TB of disk I/O.

This inefficiency has been well known for years, but there is little reward or funding for recreating existing tools. The pipeline works well, if slowly, and rewriting all the tools by hand would be an enormous task. Coding agents have changed this: the cost of reimplementation has dropped to a few days, making it worth collapsing the redundant passes into one and recovering the wasted compute, provided the numbers downstream analyses rely on did not change. By targeting a rewrite rather than a new tool, the difficulties of adoption and trust could largely be sidestepped while performance improved radically.

In the months since RustQC’s release, I have been involved in several more rewrites: Trim Galore (quality- and adapter-trimming), which has been led by its original author Felix Krueger; FastQC (raw sequence QC), where the original author Simon Andrews did not want the canonical project replaced by a Rust rewrite, so I am instead trying to make the rewrite redundant through upstream contributions; and, finally, the aligner itself, STAR, described in another case study in this paper.

### Agentic intervention

The term “rewrite” is open to interpretation. Here I steered the coding agent to combine multiple tools into a single-pass binary: an architectural redesign aiming for maximum performance while keeping outputs intact. This puts RustQC at the “new tool” end of the scale, rather than maintenance or a targeted patch. Agents were given the source code for the original tools and their dependencies and instructed to fold their functionality into RustQC. In several places underlying C libraries were used to keep outputs identical. The agents had free rein over the codebase to explore, prototype, benchmark and plan.

RustQC was not my first attempt at such a rewrite. Earlier tries in 2025 were short-lived and unsuccessful; it took the latest advances in both models and agentic harnesses for a project this complex and long-lived to become viable.

Trim Galore, FastQC-Rust and ruSTAR are more traditional rewrites: faithful 1:1 reimplementations aiming for exact emulation of the original, including its usage experience, rather than redesign.

### Human role

My role in RustQC was domain expert, architect and validator. I chose what to build and what to leave out, for example scoping RustQC to the minimal feature-set needed to replicate the nf-core/rnaseq QC subworkflow. Perhaps most importantly, I built the validation infrastructure. In practice this meant steering the model over many hours, deciding priorities and tradeoffs, and correcting it when it made false assertions. This mattered most in the later stages: the initial work took days, but refining the code to output parity took a further month, during which heavy human oversight was essential to ensure the tool reproduced the originals faithfully across many edge cases.

I did not review the implementation line by line; instead, I ensured correctness by comparing outputs against the original tools.

### Validation and evidence

Agents are eloquent, convincing, and confidently wrong in ways that are easy to miss, so I never let the model decide whether its own output was correct. I always used a rigorous test harness. I started with minimal single-tool benchmarks. As more tools were added I built a Nextflow benchmarking pipeline with unit tests. Eventually I used the nf-core/rnaseq pipeline itself: run it with the original tools, run it again with RustQC, and compare. Nextflow gives deterministic execution, with cached intermediates. nf-test adds file snapshot assertions, so every change was checked against a fixed reference. Any divergence in a value, column, header or filename was clearly surfaced at once.

I validated against real public sequencing data across several organisms and library preparations, because real data carries error and quality profiles that are difficult to reproduce with synthetic data. Volume mattered too: many edge cases only surfaced at realistic scale, and minimal datasets were not sufficient.

### Outcome

On a large paired-end human dataset of around 186 million reads run through nf-core/rnaseq on AWS, the original tools take about 15 hours 34 minutes of sequential runtime, of which RSeQC’s TIN alone is roughly 9 hours 45 minutes. RustQC finished the same work in 14 minutes 54 seconds, more than 60x faster, with disk I/O down from 2.5TB to 0.1TB. The output is numerically equivalent and MultiQC-compatible, so switching a pipeline over is close to a one-line configuration change and is reversible.

Other rewrites also gave strong speed improvements, with Trim Galore 7x faster and FastQC-Rust 3x faster. Through continued agentic work with my contributions to the upstream Java implementation of FastQC, I have since managed to get the same 3x speed-up in the original codebase.

rewrites.bio prompted an extended discussion across social media and bioinformatics boards, with comment threads running into the hundreds. Many people supported its aims, and a number of people have begun to refer to its principles in their own rewrites.

### Obstacles and failure modes

Agents were good at the things that usually take the most time if done manually: porting algorithms between languages, producing large amounts of correct Rust code despite my inexperience, scaffolding tests and documentation, and, when asked, profiling and improving their own code. However, their weakness was the one that perhaps matters most in scientific software: judging their own correctness. The failures were almost never crashes; they were typically small numerical and formatting divergences. The agent was nearly always aware of these differences but would drift from the original prompts in an attempt to complete the task and declare them “scientifically valid” or “acceptable”. This is why the external harness was necessary. I also had to coerce the agent to make errors and unsupported functionality fail loudly, rather than silently return the wrong answer.

### Lessons and reflections

The main lesson for me was that fast code has become cheap while correctness, attribution and trust have not. I found it helpful to think big, but work small: RustQC showed that huge performance wins can come from rethinking the architecture, reducing I/O, skipping intermediate steps and combining tasks, but the work itself only succeeded when it proceeded in small steps, each validated against the original before the next began. I think that there is a definite problem with ongoing maintenance costs and rewritten tool stewardship, which I do not think the field has answered. A wave of AI-assisted rewrites has already begun, and if they diverge in behaviour they will fragment the community. It’s critical that results from different labs and different times (and potentially different rewrites) can still be used together.

Overall I am extremely optimistic about the agentic future of bioinformatics. Coding has been the bottleneck on what most of us can achieve in a day, and technical debt has been tolerated because we had little choice. Agents change that and if used responsibly, the shift is firmly for the better.

### Artifacts

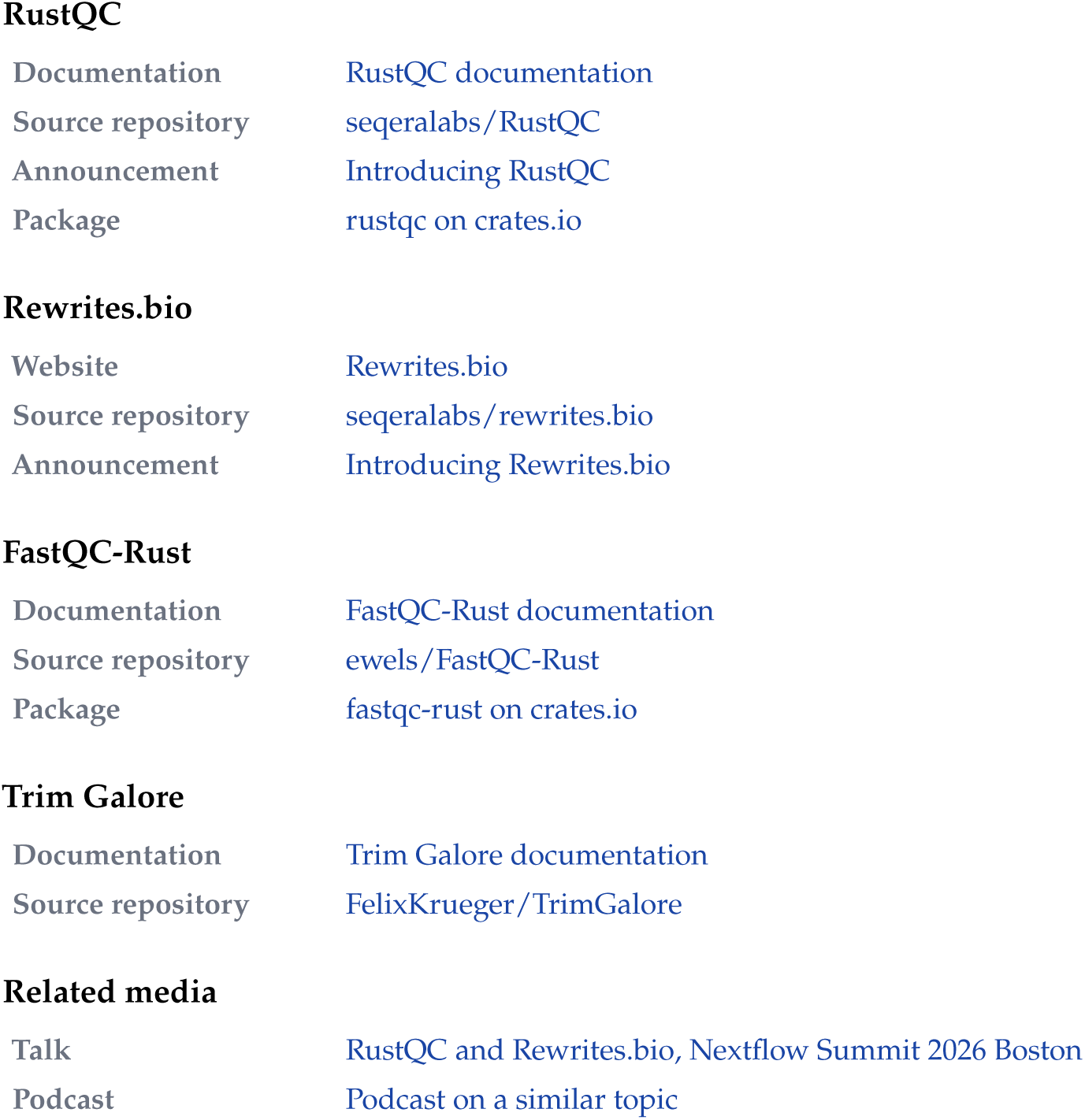

RustQC
|  |  |
| --- | --- |
| Documentation | RustQC documentation |
| Source repository | seqeralabs/RustQC |
| Announcement | Introducing RustQC |
| Package | rustqc on crates.io |
| Rewrites.bio |  |
| Website | Rewrites.bio |
| Source repository | seqeralabs/rewrites.bio |
| Announcement | Introducing Rewrites.bio |
| FastQC-Rust |  |
| Documentation | FastQC-Rust documentation |
| Source repository | ewels/FastQC-Rust |
| Package | fastqc-rust on crates.io |
| Trim Galore |  |
| Documentation | Trim Galore documentation |
| Source repository | FelixKrueger/TrimGalore |
| Related media |  |
| Talk | RustQC and Rewrites.bio, Nextflow Summit 2026 Boston |
| Podcast | Podcast on a similar topic |

### References

1. Ewels PA, Peltzer A, Fillinger S, Patel H, Alneberg J, Wilm A, Garcia MU, Di Tommaso P, Nahnsen S. The nf-core framework for community-curated bioinformatics pipelines. *Nature Biotechnology.* 2020;38:276–278.

### CASE STUDY D

### Agent-driven development of a GPU-native engine for synthetic genome generation

**Mamad Ahangari^1^, Varun Goyal^1^, Hassan Masoudi^1^**

^1^ MinosAI

### Summary

Mutation detection is a core task in human genomics. The tools that perform this work, known as variant callers, need to be evaluated against genomes with known ground-truth mutations. This creates a bottleneck because only a small number of human genomes have sufficiently complete ground truth to support rigorous evaluation.

The usual workaround is spike-in data generation, where real genome sequence data is edited by inserting mutations at known locations, then using those edits as the ground-truth answer key. BamSurgeon^1,2^ is the standard tool for this edit-and-label step, and it has supported a decade of method development and benchmarking. Its design is generally reliable but artifact-prone and computationally expensive. For each mutation, the tool launches a chain of general-purpose genomics tools, including bwa-mem, Picard, and samtools. This makes it accurate enough to trust the injected variants, but slow to scale. It also re-aligns every edited read, which leaves a detectable local signal at the sites it modifies. These scaling and artifact issues are not unique to BamSurgeon, as other less widely used methods also suffer from the same limitations.^3,4^

This case report describes how we rebuilt the existing spike-in workflow pipelines with AI coding agents to develop HelixForge, a GPU-native engine for scalable synthetic genome generation at population scale. Over a period of about a month, we used OpenAI GPT-5.5 Pro API and Codex to identify and harden the artifact-prone and bottlenecked steps in BamSurgeon, and then replaced them entirely with an updated workflow that uses a custom htslib and CUDA C++ engine on an NVIDIA H200 GPU and, on the matched benchmark reported here, outperformed BamSurgeon in both speed and measured mutation-frequency accuracy. On a like-for-like benchmark, HelixForge is about 60 times faster end-to-end and about 99 times faster in the genome editing step. On this benchmark, it produces fewer detectable artifacts and nearly eliminates the measured re-alignment signature, cuts mutation-frequency error by more than half, and reproduces every requested mutation in the benchmark. The tool can spike-in single-base substitutions (SNP), short insertions and deletions (INDELs), as well as structural, copy-number, and cancer-type mutations. We show that beyond the significant speedup and reduction in artifact signatures, this workflow allowed us to use agents to identify and fix real defects in a trusted existing tool widely used in genomics by establishing a strong validation harness that pushed beyond the original design of existing state-of-the-art spike-in tools.

### Scientific context

There are two broad ways to create synthetic test data for mutation detection in genomics. One is to simulate sequencing reads from scratch. This approach is relatively fast, but it creates a distribution shift because simulated reads only approximate the error profile of a real sequencer as variant callers can behave differently on simulated reads than on real data.^3^ Another approach is to use Spike-in methods which avoid that problem by editing real reads from sequence data instead. Spike-ins keep the noise, coverage, and machine-specific quirks of the original sequencing run, while letting you choose the mutations and labels by injecting them at precise locations in the genome. Such spike-in approaches are generally more realistic than entirely simulated genomes, but they are still prone to artifacts and are significantly slower and more computationally expensive.

### What BamSurgeon is, and why it is slow and artifact-prone

BamSurgeon is the most widely used example of spike-in approaches for generating synthetic genomes. The tool is best understood as an orchestration layer for coordinating multiple external bioinformatics tools, repeatedly rewriting, re-aligning, sorting, and repairing reads around each injected mutation. This design makes it useful for realistic synthetic truth construction, but very prone to artifacts and difficult to scale beyond small targeted regions.

In our benchmarks, the primary task was injection of 100 single-base substitutions plus 5 insertions or deletions in a 10 Mb region. BamSurgeon took 1,610 seconds on average, and 1,557 seconds of that was the editing step alone. Using this workflow, we identified another limitation of BamSurgeon’s orchestration pipeline beyond its high computational cost. BamSurgeon edits reads and then re-aligns them to the reference genome. As a result, the modified reads can show a mapping-quality distribution that differs from that of nearby, unmodified reads, which creates a detectable signal around the spike-in site.^1^

Mapping quality reflects the aligner’s confidence that a read has been placed correctly in the right location of the genome. In real sequencing data, these values typically vary across reads. By contrast, re-aligned spike-in reads can cluster in a way that makes them distinguishable from the surrounding background. This issue is not necessarily specific to BamSurgeon though. Any method that modifies reads and then processes them again through an alignment pipeline may introduce these signatures. Previous studies have noted that existing simulation tools often provide limited control over the biological and technical properties of their outputs which remains a broader challenge in synthetic genome generation.^3,4^

### Development of a scalable spike-in engine

We developed HelixForge through an agent-assisted workflow. The scientific goals, development sequence, evaluation criteria, and acceptable failure modes were all defined by our research team, while AI coding agents were used for much of the implementation and iterative debugging of the work.

Instead of attempting to build the entire GPU-native pipeline at once, we first used the agents to improve existing BamSurgeon workflow and characterized the ways in which its output could appear unrealistic or retain inconsistent metadata. These inconsistencies and issues included read groups being overwritten by Picard, fixed rather than variable allele fractions, a general lack of sequence context, uniform INDEL along with outdated consistency tags in the metadata and the absence of explicit strand-balance and coverage checks. This initial work to identify failure modes produced a stable CPU reference that could be used to directly evaluate the GPU-based model against it using controlled A/B comparisons. Based on the runtime profiling and artifact analysis, we then replaced the pipeline stages that contributed most to the computational bottleneck with a GPU-native implementation and nearly eliminated the measured re-alignment fingerprint.

In short, we defined the scientific objective, failure modes, validation criteria, and build order, while AI agents handled much of the implementation, CUDA iteration, source review, and validation harness development.

### Validation of the workflow

Every comparison in this report uses matched inputs for CPU-bound BamSurgeon and GPU-native HelixForge. Both tools were run on the same donor genome, HG005 from Genome in a Bottle (GIAB) consortium. The same 10 Mb region of chromosome 20 with the same set of mutations and random seeds were used for all comparisons throughout.

This allowed us to make the comparison close to an ablation study where the main intended difference is the mutation-injection engine. We report per-run averages and present this as a focused engineering measurement, not an exhaustive cross-donor benchmarking analysis.

To isolate the components we replaced in HelixForge, we separate wall-clock time into the editing step and the surrounding preparation and finalization work. On the editing step, BamSurgeon averaged 1,556.9 seconds, while HelixForge averaged 15.8 seconds for the same task, making the GPU-native path about 98.6x faster. End-to-end, BamSurgeon averaged 1,609.6 seconds compared with 27.0 seconds for HelixForge, a 59.6x speedup. The editing-step ratio is the cleaner comparison because it isolates the replaced component from finalization work shared by both paths.

The injected mutations also improved in quality with the GPU-native path, despite the large reduction in runtime. We measured the synthetic fingerprint as the fraction of edited reads carrying the tell-tale mapping-quality pattern introduced by re-alignment. BamSurgeon was essentially 1.0 on this measure, with about 1,971 fingerprinted reads per run. In contrast, HelixForge was close to 0, with about 0.4 such reads per run. Mutation-frequency accuracy also improved, with the average error falling from 0.076 to 0.034. For insertions and deletions, the correlation between requested and observed frequency increased from 0.80 to 0.99. HelixForge also reproduced every requested spike-in mutation, compared with a 99.7% confirmation rate for BamSurgeon. In short, the new pipeline made the process both faster and more faithful across every measure we tested (Figure D1).

**Figure D1.**
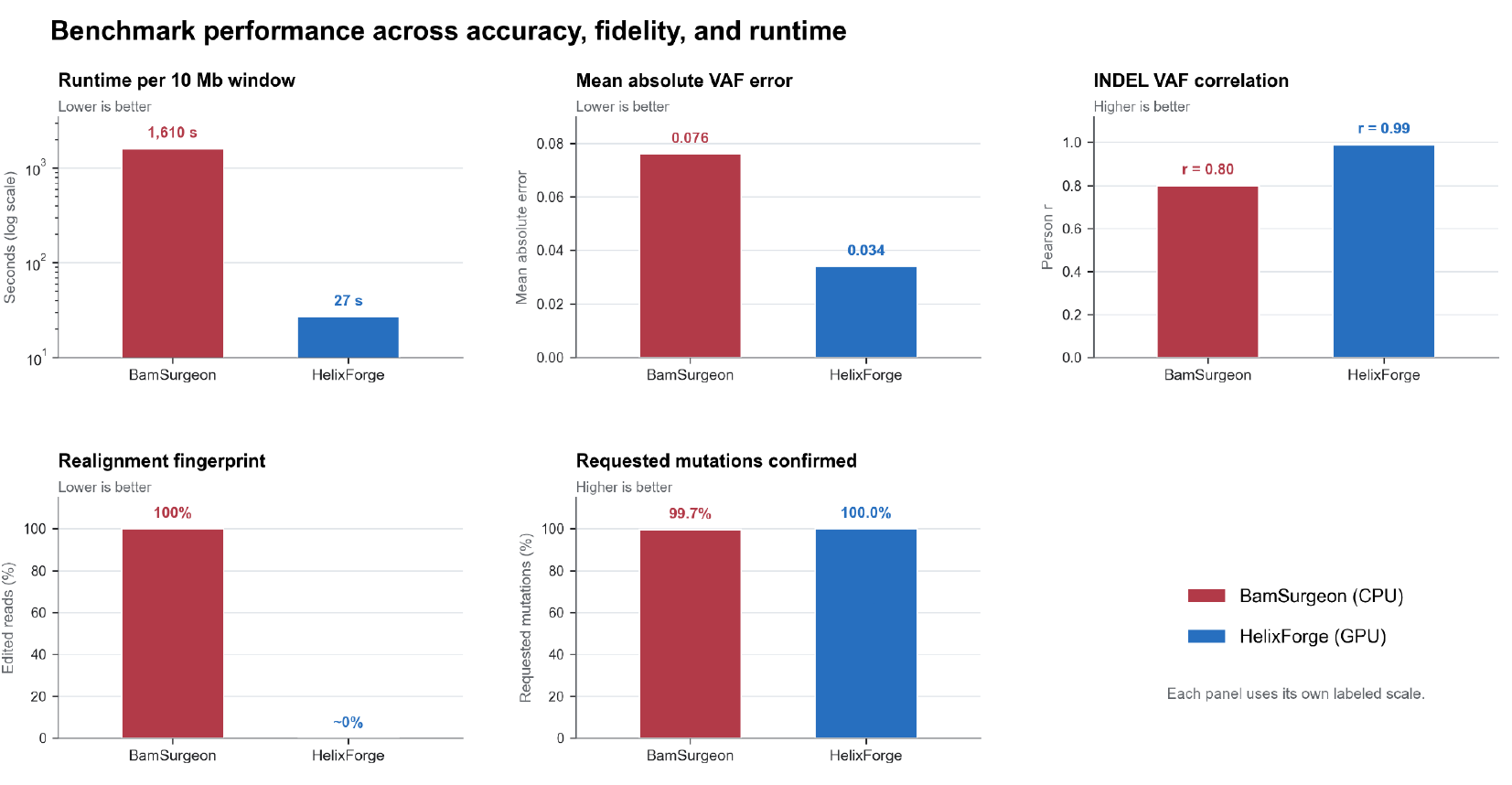
BamSurgeon and HelixForge on an identical task using the same donor, region, mutation sets, and seeds. Each metric is shown on its own scale, with the better value on the right. HelixForge is faster and less artifact-prone on every measured axis.

The accuracy gap is clearest at individual mutation sites. When the requested spike-in mutation frequency is plotted against observed frequency for each confirmed mutation, HelixForge stays close to the line of perfect agreement, whereas BamSurgeon shows systematically wider scatter, with several insertions and deletions collapsing toward zero observed support (Figure D2).

**Figure D2.**
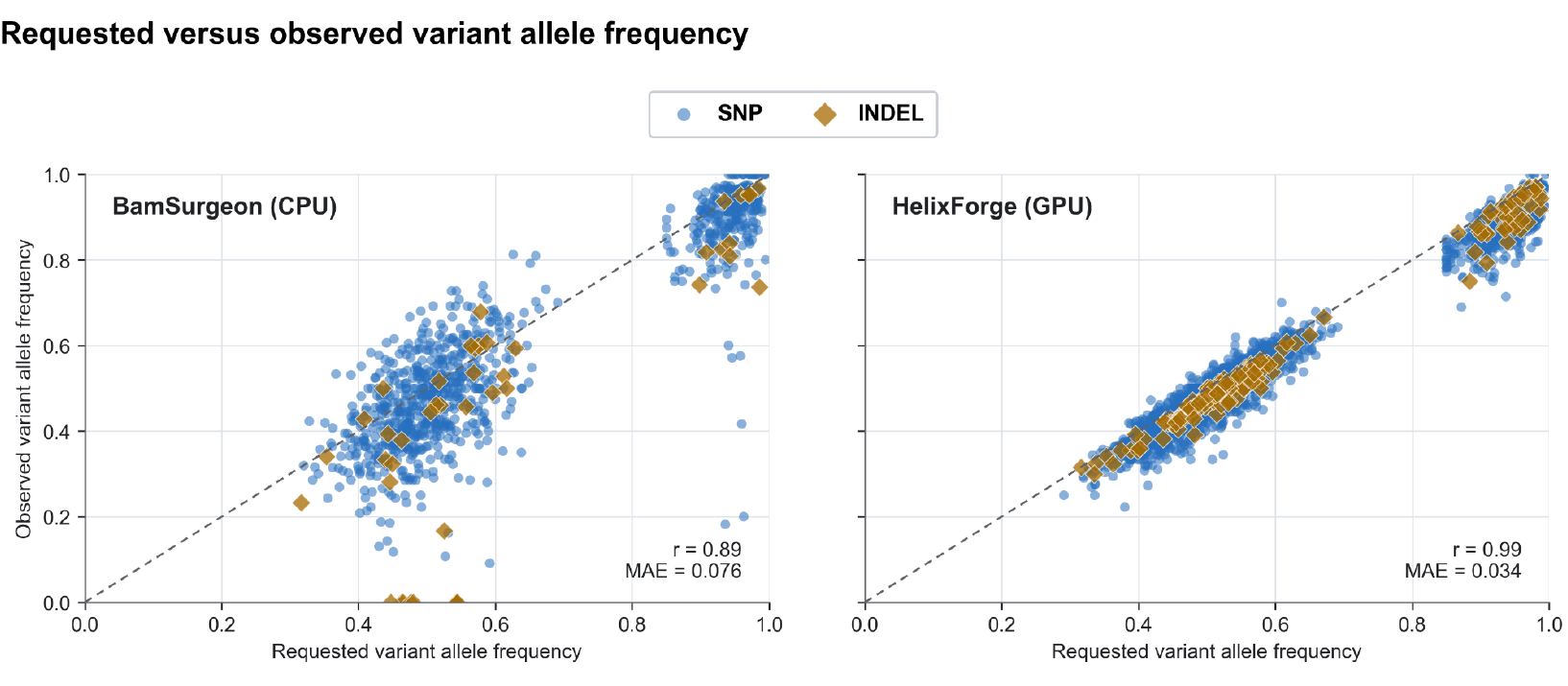
Requested versus observed variant frequency for every confirmed mutation on the HG005 chromosome 20 task. The dashed line indicates perfect agreement. HelixForge on the right tracks it closely. BamSurgeon on the left scatters, with the largest errors on insertions and deletions.

### Parallelization of the workflow with agents

Our goal in this study was not to create a single synthetic genome example but to build the infrastructure needed for a growing library of synthetic genomes at scale. While a one-off spike-in genome can support a local test, large-scale benchmarking requires many independently generated synthetic genomes with unique spike-in properties. We therefore used the agent-driven development loop to design the workflow around clean parallelism from the start, ensuring that separate files shared no state and could be generated independently across multiple GPUs.

This parallelism is what turns the GPU-native engine from a faster tool into a scalable data-generation system. On eight H200 GPUs in one machine, we show that throughput scales almost linearly, reaching an effective

3.6 seconds per 10Mb genomic file. At that rate, a synthetic whole genome, modeled by tiling a 3.1 Gb genome into approximately 310 ten-megabase windows, is projected to require about 2.3 hours on one H200 or approximately 19 minutes when the windows are distributed across eight H200s. The corresponding projected runtime for the BamSurgeon CPU path is approximately 5.8 days per genome (Figure D3).

At a larger scale, a cohort of approximately 3,200 synthetic whole genomes at 30x coverage would require about ten months on one H200 or about six weeks on eight H200s operating in parallel. The equivalent BamSurgeon CPU workload would require approximately 51 years on a single worker (Figure D4). This scale is only useful if the synthetic data remains difficult to distinguish from real sequencing data, which is why HelixForge uses in-place editing rather than BamSurgeon’s native re-alignment-based spike-in generation, which leaves multiple detectable artifacts in the generated genomes.

**Figure D3.**
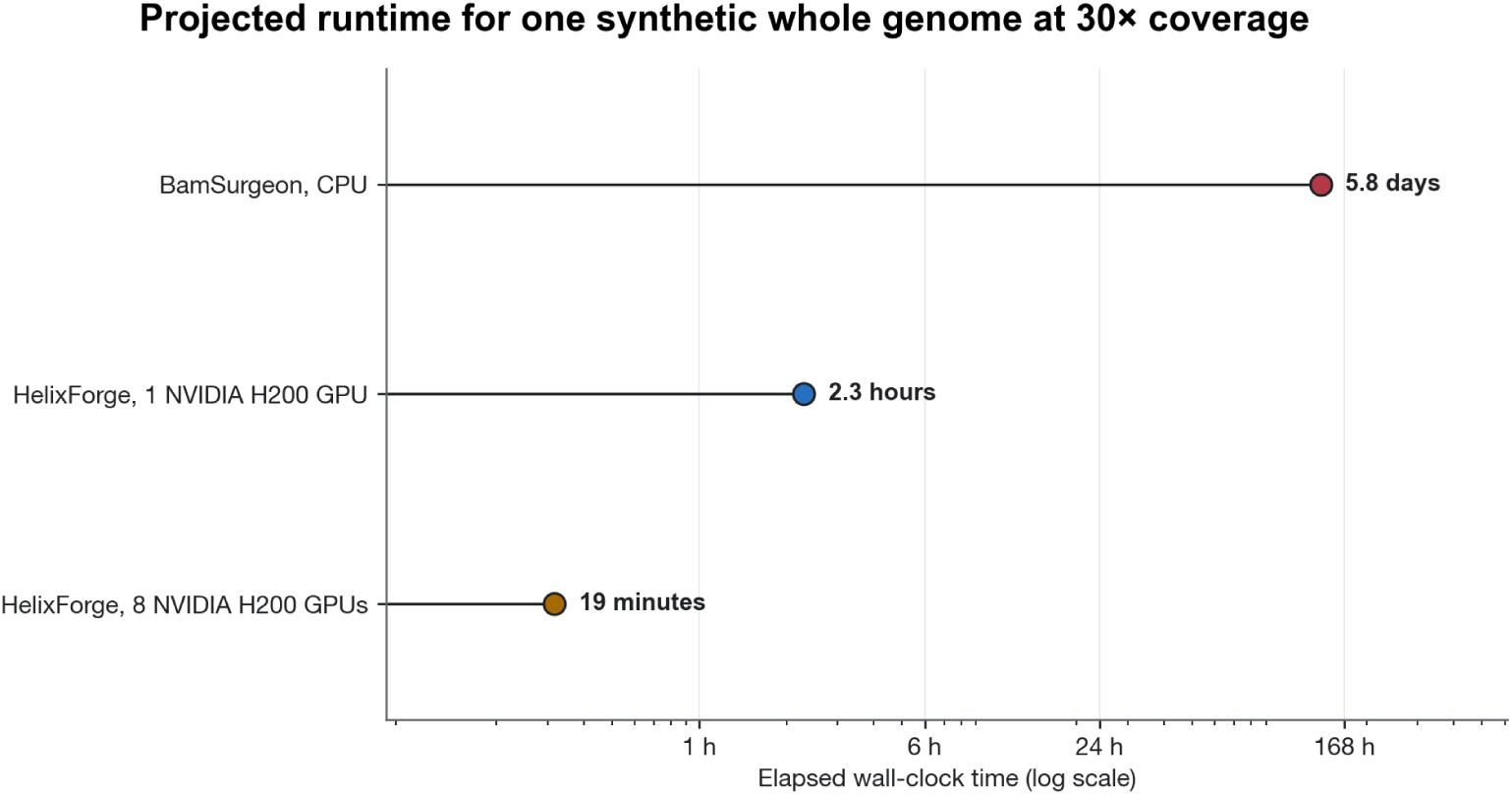
Projected wall-clock time to generate one synthetic whole genome at 30*×* coverage. Per-window measurements were 1,609.6 seconds for the BamSurgeon CPU path, 27.0 seconds on 1 NVIDIA H200 GPU, and approximately 3.6 seconds across 8 NVIDIA H200 GPUs; scaling these values across approximately 310 10-Mb windows gives projected whole-genome times of 5.8 days, 2.3 hours, and 19 minutes, respectively.

**Figure D4.**
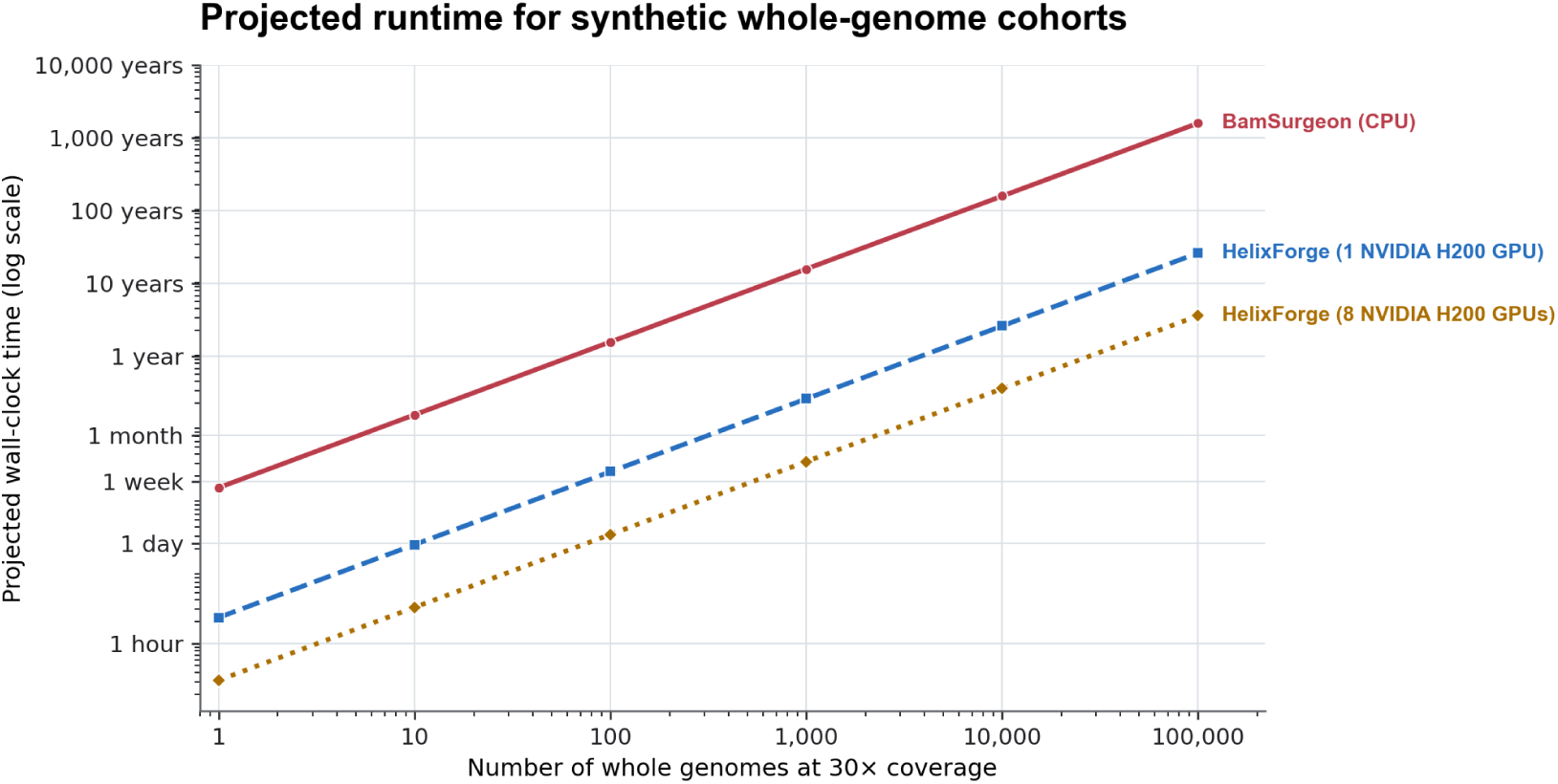
Wall-clock time to generate cohorts of synthetic whole genomes at 30x coverage using the BamSurgeon CPU path, 1 NVIDIA H200 GPU, or 8 NVIDIA H200 GPUs using HelixForge.

### The human role

Although the agents handled much of the implementation, expert human judgment remained the most crucial part of the entire project. One of the main responsibilities was deciding the order of tasks and what to work on at each step. Agents performed well when given a clearly defined task, but they were less reliable at setting the development sequence themselves. We therefore chose to stabilize the CPU reference workflow first before attempting to build the GPU engine as described above. Without that direction, the work often shifted toward local improvements or unnecessary complexity rather than the most direct route to a validated system with clear objectives.

Expert human review was also required at each validation checkpoint. We did not treat an agent’s report that a component was complete, or that its tests had all passed as sufficient evidence on its own. Each major change was followed by same-seed comparisons and additional checks selected and reviewed by the research team. This allowed us to measure performance gains without losing biological accuracy or file-level consistency.

The third area of human involvement was scientific realism. An agent can produce code that compiles and generates a formally valid genome sequence file, but that does not guarantee that the output will appear realistic to an experienced geneticist. Several of the auditing steps that are included in HelixForge were added because intermediate outputs, both generated by BamSurgeon, and further by the initial versions of HelixForge, were technically valid yet contained patterns that were recognizably synthetic by a trained geneticist. The final workflow therefore used agents for implementation speed and breadth of work, while leaving development priorities, validation standards, and scientific realism and interpretation under expert human control.

### Obstacles and lessons

We consistently observed that agents were most effective when their task specifications were narrow and the tests were fully objective. Once the same-seed comparison framework was in place, the agents could modify GPU routines, translate logic between different coding languages, and iterate on implementation details until the results matched our desired speed and output.

The agents were less reliable when the source of an error was more ambiguous. For example, in one instance, an early strand-balance audit produced a false positive because of the downsampling procedure. The agents responded by modifying the GPU implementation, even though the problem was in the auditing step itself. This reinforced the need to determine whether a failing test reflects an implementation error or an incorrect assumption in the first place in the test.

Agents also tended to bypass difficult edge cases unless those cases were made explicit in the specification in advance. For example, early work on INDEL injection did not fully handle overlapping edits. The resulting genomes appeared plausible but were incorrect. We addressed this sort of issue by requiring the pipeline to stop with a clear error whenever an edit could not be handled safely, rather than silently producing uncertain outputs.

Finally, we learned that broad instructions often led to unnecessary changes outside the intended scope of the task. This is particularly risky in pipelines where reproducibility depends on the exact order of operations, random-number generation and sampling behavior in the code. For that reason, we ensured that later tasks were narrowly scoped and changes were reviewed against the reference implementation before being accepted into the pipeline.

The single most useful artifact in our study was the same-seed comparison harness. Once it existed, nearly every later step became an iteration against a stable reference. The second unlock was keeping durable, human-readable design notes inside the repository. These notes gave the agents continuity across sessions and model versions, while keeping the scientific assumptions visible to the human reviewer.

Overall, the agents were an unambiguous net positive when used inside a tight validation loop with human-controlled framing and acceptance criteria. While they were less reliable when asked to make open-ended scientific judgment, we learned that having a harness significantly helped with open-ended experimentation and feature development in the context of genomic tool development.

### Artifacts

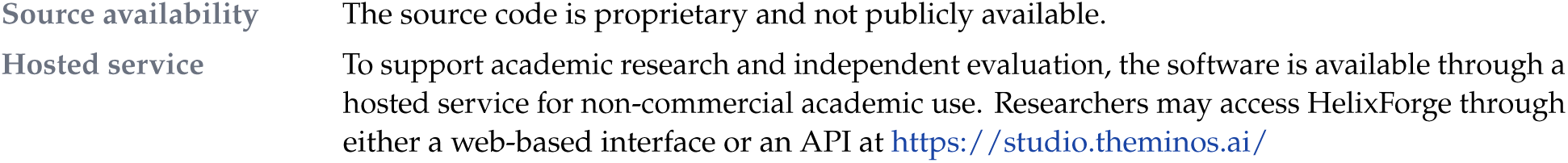

### References

1. Ewing AD, Houlahan KE, Hu Y, et al. Combining tumor genome simulation with crowdsourcing to benchmark somatic single-nucleotide-variant detection. *Nature Methods.* 2015;12:623–630. doi:10.1038/nmeth.3407.

2. Lee AY, Ewing AD, Ellrott K, et al. Combining accurate tumor genome simulation with crowdsourcing to benchmark somatic structural variant detection. *Genome Biology.* 2018;19:188. doi:10.1186/s13059-018-1539-5.

3. Milhaven M, Pfeifer SP. Performance evaluation of six popular short-read simulators. *Heredity.* 2023;130:55–63. doi:10.1038/s41437-022-00577-3.

4. Longhin F, Baruzzo G, Hazizaj E, et al. MOV&RSim: computational modelling of cancer-specific variants and sequencing reads characteristics for realistic tumoral sample simulation. *BMC Bioinformatics.* 2025;26:287. doi:10.1186/s12859-025-06292-0.

### CASE STUDY E

### Agentic optimization of a genome assembly library

**Suyash Shringarpure^1^**

^1^ OpenAI

### Summary

Hifiasm is a widely used genome assembler for PacBio HiFi reads. We explored whether an automated, LLM-driven optimization process could improve its performance while preserving assembly quality. We used smaller synthetic datasets for quicker evaluation and real human data for final testing. On a held-out 200 Mb synthetic benchmark, the best optimized implementation from GPT-5.5 reduced runtime by 25.1%, while satisfying predefined assembly-ordering quality thresholds. The improvement also transferred to human long-read data from the Human Pangenome Project, where runtime decreased by 14.7%.

### Scientific context

Hifiasm was developed by Cheng et al. in 2021 for haplotype-resolved de novo genome assembly. Many genome assembly algorithms collapse heterozygous alleles into one consensus copy or fail to separate haplotypes. Hifiasm uses long high-fidelity sequence reads to summarize haplotype information in a phased assembly graph. It is the state-of-the-art approach for de novo assembly, with modifications to account for newer sequencing technologies with different quality and error profiles.

### Problem or opportunity

Hifiasm can assemble a full human genome in 6–8 hours. We explored whether agentic optimization could be used to reduce this runtime. An expensive step in the hifiasm algorithm is the all-vs-all read overlap alignment. We hypothesized that this could be an area where runtime improvements were possible. Hifiasm is a large piece of software, with 178,086 lines of code (LoC) across 59 tracked C/C++ source/header files. Therefore, this offers an opportunity to test whether coding agents can reliably modify large codebases.

### Agentic intervention

We first built a reproducible benchmark harness covering multiple dataset sizes and established separate development and held-out evaluation datasets. Profiling showed that read correction, edit-distance computation, trace generation, and overlap chaining dominated runtime. Optimization efforts therefore focused broadly on reducing repeated work in these hot paths, improving memory-access patterns, and introducing efficient fast paths for common cases. GPT-5.5 was prompted by providing profiling results and instructions to optimize runtime while keeping quality unaffected.

Candidate implementations were tested on a small 12 Mb development dataset to catch compilation failures, performance regressions, and quality problems early. Promising candidates were then evaluated from a clean source tree on the held-out 200 Mb synthetic dataset. The evaluation independently measured the unmodified baseline and optimized implementation and rejected changes that failed assembly-quality constraints.

To test whether improvements found using simulated data generalized to real reads, we also benchmarked optimized candidates on chr 20 reads from human HG02723 sequencing, using the same clean-build comparison against the unmodified source.

### Human role

The human contribution was primarily to design the problem and evaluation setup.

- Creating a development dataset and a held-out evaluation dataset, as well as a second held-out dataset.
- Designing the model prompt to include profiling results.
- Based on early attempts, revising the model prompt to prevent common failure modes.

### Validation and evidence

Since comprehensive evaluations of genome assembly quality can be complex and byte-level equivalence can be an overly rigorous standard for an optimization exercise, we used a read-ordering metric as a proxy for assembly quality. To create this metric, we used the ordering of non-contained reads from the synthetic ground truth in the baseline hifiasm assembly as our baseline metric. For every optimization candidate, an assembly was produced on the same dataset, requiring 100% concordance (within 1e-12 floating-point tolerance) with the baseline on the read-ordering metric for quality acceptance. Any candidate which did not attain this concordance was discarded.

Several candidate proposals achieved substantial improvements while meeting the assembly quality requirements. The strongest result reduced held-out synthetic runtime by approximately 205 seconds, from 816.9 seconds to 612.0 seconds, while preserving the required ordering-quality metrics.

The performance gain transferred to the human data from the Human Pangenome Project, although at a smaller magnitude. For this evaluation, we used data for chromosome 20 from HG02723. Runtime decreased from 734.8 seconds to 626.6 seconds, saving 108.2 seconds and producing a 14.7% reduction. Overlap counts differed by only 43 out of 10.77 million, suggesting that the optimized assembly was almost identical to the original hifiasm assembly.

### Outcome

This work shows that coding agents can effectively propose optimizations to large codebases. In this case, the agent autonomously identified an area for improvement, implemented a change, tested it against a development dataset and benchmarked its improvement. The agent successfully proposed and implemented a change that reduced runtime by 25% on the synthetic dataset and by 14.7% on the human dataset while preserving assembly quality. Our results suggest that agentic optimization benefits can extend along the full spectrum of scientific software from small codebases with thousands of LoC to large ones with hundreds of thousands of LoC.

### Obstacles and failure modes

Initial attempts at prompting the LLMs using only the source code and a high-level prompt revealed a number of failures. Initial optimization proposals were myopic and influenced by the small size of the development dataset, suggesting trivial improvements such as reducing the number of correction rounds, or hardcoding the size of data structures based on the small dataset. Another failure mode was to focus improvements on parts of the algorithm that only accounted for a small proportion of overall runtime. Both these failure modes were fixable by providing appropriate context, through detailed instructions for the former, and through profiling results for the latter.

### Lessons and reflections

This work demonstrates that meaningful performance improvements can be recovered from mature, specialized genomics software without weakening the quality requirements used during optimization. Profiling, representative benchmarks, and clean held-out evaluation were critical: many plausible optimizations performed well on small workloads but did not generalize, while the strongest changes produced consistent improvements at larger scale.

The human data results show that the gains were not confined to the synthetic benchmark. The runtime reduction remained substantial on human data, though it decreased from 25.1% on the held-out synthetic workload to 14.7% on the recorded human workload. This attenuation highlights the importance of testing optimized systems across data distributions rather than relying only on simulated benchmarks.

Human contribution in the optimization process was critical in designing the problem and evaluation, as well as in providing the model with suitable context to guide its proposals. Once those had been established, the optimization process was fairly hands-off, and the model proposed strong candidates even on a mature codebase. Making coding attempts easily accessible to genomics software developers could help speed up a large portion of the genomic software ecosystem and would have positive downstream effects on users of such software.

### Artifacts

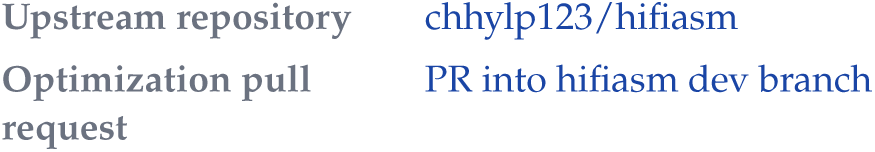

### CASE STUDY F

### Cyvcf2: Case study in maintaining and distributing software

Brent Pedersen^1^

^1^ Independent researcher

### Scientific context

One of the largest burdens in building scientific software is long-term maintenance. Post-docs and graduate students move on to new positions and staff scientists move on to other projects. There is also little funding to support long-term development of software. It is therefore common for projects to experience bit-rot, to only work on legacy systems, and to rely on outdated build pipelines.

### Problem or opportunity

Cyvcf2 (https://doi.org/10.1093/bioinformatics/btx057) was created over a decade ago to facilitate rapid reading and writing of VCF files. Since that time, there have been many Python version changes (2.7 to 3.14) and the Python build system has gone through many iterations. In addition, when *cyvcf2* was created, GitHub Actions, used for automated continuous integration and deployment, did not yet exist. The complexity of simply keeping the build-system up-to-date is evidenced by the commit history of cyvcf2 (https://github.com/brentp/cyvcf2/commits/main/) where most commits deal with dependencies and build issues. For an open-source project, this is an immense maintenance burden. This is multiplied, over the interval of software maintenance, by all projects maintained by a given developer. The aim of this project was to harden and modify the cyvcf2 build and test infrastructure which has been held together by contributors.

### Agentic intervention

Given the long history of maintenance of *cyvcf2*, much extra cruft and complexity had accrued. A coding agent with a recent training cutoff has knowledge of Python packaging best practices and can recommend and implement a modern approach to Python packaging, testing, and distribution. The existing tests make this process more stable as there is a benchmark to evaluate on.

### Obstacles and failure modes

This project was a major revamp of the build system for *cyvcf2*. We found it useful to have two separate pull-requests; one was a minor change, and the other more substantial. Only the more substantial pull request was ultimately merged. This was possible because of the efficiency of agents and allowed the *cyvcf2* maintainers to weigh the benefits of each approach. The major change used a module that was less familiar to *cyvcf2* maintainers (scikit-build-core) and so it was useful to have external agents further review the code. Those agents and human reviewers found several small checks to be added and changes that hardened the change. Combining human and agent review proved quite useful. In the end, it was human guidance that led to the specific implementation. Confidence in the implementation was possible because of the test-suite and continuous integration; without that, it would be difficult to fully evaluate the consequences of such a large change.

### Lessons and reflections

While not glamorous, this project represents a practical and useful case-study. This change will reduce the maintenance burden of the software going forward. Long-term maintenance is crucial for the bioinformatics infrastructure and can be especially burdensome for scripting languages and for projects that have existed (or will exist) for more than a decade. The tedious changes here required an up-to-date and thorough knowledge of the Python packaging ecosystem that scientific developers often do not have time to cultivate. Having thorough, human-written tests facilitates this process.

### Opinionated best practices

The rise of agentic coding changes things for software in genomics. Many tools are written for bioinformaticians or even adept clinicians; adding options to a command-line tool is a way to make knobs accessible to those users without exposing too much complexity. Now, agents can use those options, but, since agents are so good at coding, it can be even more powerful to expose a scripting interface to give nearly infinite customizability. This can allow the user to do things that the implementor didn’t even consider. The agent can write, for example, JavaScript expressions to filter genetic variants as in slivar, and a custom GPT or a skill can tell an agent exactly how to utilize it. The result can be a script that, once written, is simple enough for a human user to modify and tweak as needed, or to iterate on with the agent. So far, exposing a scripting interface in this way is an underutilized avenue in agentic coding.

We are still in the early days of agentic coding and best practices will surely change. It seems nearly certain that at some point, we won’t read the code, just as now, we don’t read the machine code of our higher-level source code. As of mid-2026, several things remain for a human to do. The most important of these is *care* for the software process. An agent will now readily one-shot an entire codebase, but without human involvement and care, there is no learning. Even with a careful design, software evolves during implementation along with the understanding of the (human) implementor. If we leave ourselves out of the loop, this does not happen. In addition, it is possible to move so fast with agentic coding that it’s disappointing to read all or even some of the code. Without care, the code will pass tests and may look impressive in benchmarks but is a net negative on the community.

### Artifacts

|  |  |
| --- | --- |
| Source repository | brentp/cyvcf2 |
| Merged pull request | PR #327: Migrate native build to scikit-build-core |
| Release tag | v0.33.0 |

### CASE STUDY G

### bayesm rewrite and extensions

**Andrew Bai^1^, Andrew Ho^2^**

^1^ Booth School of Business, University of Chicago

^2^ OpenAI

### Summary

This case study describes a Rust rewrite of bayesm using Codex agents, along with two extensions added after the base rewrite. bayesm is a widely used R package for Bayesian hierarchical models, particularly in marketing and choice modeling, making it a natural target for a rewrite. Using GPT-5.2, we rewrote bayesm in Rust as bayesm-rs, achieving feature parity with the original library alongside 2*×*–20*×* speedups on reference workloads across different functions. Across the evaluated base-rewrite workloads, bayesm-rs met the stated population-mean agreement criterion while running at least 2.31 times faster, depending on chain length and threading. These results are reported in the rewrite-validation section of this case study. The first extension is a rewrite of bayesm.HART^1^. The second adds fixed-trajectory Hamiltonian Monte Carlo (HMC) and No-U-Turn Sampler (NUTS) sampling modes^2^ to the standard bayesm functions where appropriate. We evaluated the extensions in two ways. The HMC and NUTS samplers were compared with the rewrite’s standard samplers for population-mean agreement and sampling speed. The HART rewrite was compared with the original bayesm.HART on the bank-data example from its documentation. Neither extension was correct on the first pass; later sections detail the defects and discuss lessons for practitioners implementing their own rewrites of statistical packages.

### Scientific context

The bayesm package (https://cran.r-project.org/web/packages/bayesm/index.html) is typically used in economics to fit Bayesian models, such as multinomial logit (MNL) or multinomial probit models. It was first published on May 20, 2005, by Peter Rossi, who has maintained the package since then. It is written in a mixture of R and C++. We investigated whether frontier large language models (LLMs) could execute end-to-end rewrites of existing academic libraries into higher-performance implementations and identified bayesm as a promising candidate.

This project is mainly a Rust reimplementation of the standard CRAN 3.1.7 bayesm samplers^3^. The rewrite covers the hierarchical multinomial logit finite mixture (rhierMnlRwMixture), hierarchical negative binomial (rhierNegbinRw), multivariate probit (rmvpGibbs), hierarchical binary logit (rhierBinLogit), and Dirichlet-process MNL (rhierMnlDP), among others.

### Problem or opportunity

Initially, we contemplated a variety of different approaches to this rewrite project. For example, we could have upstreamed algorithmic improvements directly to the original bayesm repository. Ultimately, we decided on our current approach—a complete, separate rewrite—because it appeared that the original library was in a very stable state and because bayesm is a fairly self-contained library which is not used upstream in a large variety of other packages. From an ownership perspective, we did not want to saddle the original author (Peter Rossi) with the task of maintaining totally foreign code, and a complete rewrite gave us the freedom to be significantly more aggressive with performance optimizations.

### Type of agent use

We prompted GPT-5.2 to execute a complete rewrite of bayesm into Rust with fairly minimal guidance. The initial prompt gave some brief instructions to check for exact parity (or statistical parity if not possible) using a suite of synthetic tests and to keep optimizations in mind. The model was then prompted with “continue” until it reported completion of the rewrite. Afterward, the model was prompted to identify, test, and implement optimizations, using a remote server for benchmarking purposes; again, fairly minimal guidance was given to the model for most of this process, with most continuation achieved by simply telling the model to “continue.”

The first extension adds a Python-facing path for nonlinear heterogeneity to three wrappers: rhierMnlRwMixture, rhierLinearMixture, and rhierNegbinRw, based on the bayesm.HART package. As opposed to standard bayesm, bayesm.HART models respondent coefficients as a tree-based function of their characteristics rather than a linear one. See https://thomaswiemann.com/bayesm.HART/ for the original implementation.

The second extension adds a shared HMC sampling option to the canonical bayesm functions alongside the standard Gibbs, Metropolis-Hastings, or hybrid routines. The original bayesm does not feature HMC as a sampling option, making this a natural extension. The engine supports two modes: fixed-trajectory HMC, in which the number of gradient steps per draw is specified manually, and NUTS, which chooses the trajectory length automatically for each draw and is the default in Stan. The HMC/NUTS option was added to rmnlIndepMetrop, rnegbinRw, rhierMnlRwMixture, rhierNegbinRw, and rhierMnlDP.

The performance improvements in bayesm-rs arose from several factors. First, GPT-5.2 identified algorithmic improvements; for example, it replaced a direct matrix inversion with Cholesky-based solves. Second, high one-time setup costs in the original bayesm implementation were eliminated in the first iteration of bayesm-rs. Third, improved memory handling, including reductions in repeated temporary allocations, reduced runtime within *hot* loops. Finally, the Rcpp bindings connecting the R interface to the underlying C++ code introduced substantial overhead. Restructuring these bindings and retaining objects inside the Rust functions for as long as necessary reduced the runtime of the R and Python bindings for bayesm-rs. Figure G1 shows the cumulative sources of speedup for rhierMnlRwMixture, evaluated through a series of code changes that simulated the historical evolution of the rewritten code.

### Human role

Achieving feature parity with the original bayesm package was relatively straightforward; because the functions are mostly self-contained, GPT-5.2 was able to progress systematically through the suite of bayesm functions and port them directly to Rust. Essentially no human intervention was required to reach feature parity, verified through numerical or statistical equivalence of outputs given identical inputs.

The work here also serves as a concise illustration of where human effort is most valuable in agentic code rewrites. The choice of original library (bayesm) and the decision to use replication of external datasets (Dubé demand estimation) as a check on rewrite validity ultimately originated from human prompting rather than from the model’s own work.

### Validation and evidence

#### Rewrite validation against CRAN bayesm

The primary validation used rhierMnlRwMixture, the package’s workhorse hierarchical choice model. Because the samplers are stochastic, exact draw-for-draw agreement is not expected. We assessed population-mean agreement by scaling each population-mean difference by the corresponding reference posterior standard deviation (SD); any scaled gap above 0.25 was considered a failure. This population-mean agreement criterion was used throughout the case study. The tables report effective sample size (ESS) per second, rank-normalized split *R̂*, and bulk and tail ESS. The convergence diagnostics follow Vehtari et al.^4^

**Figure G1.**
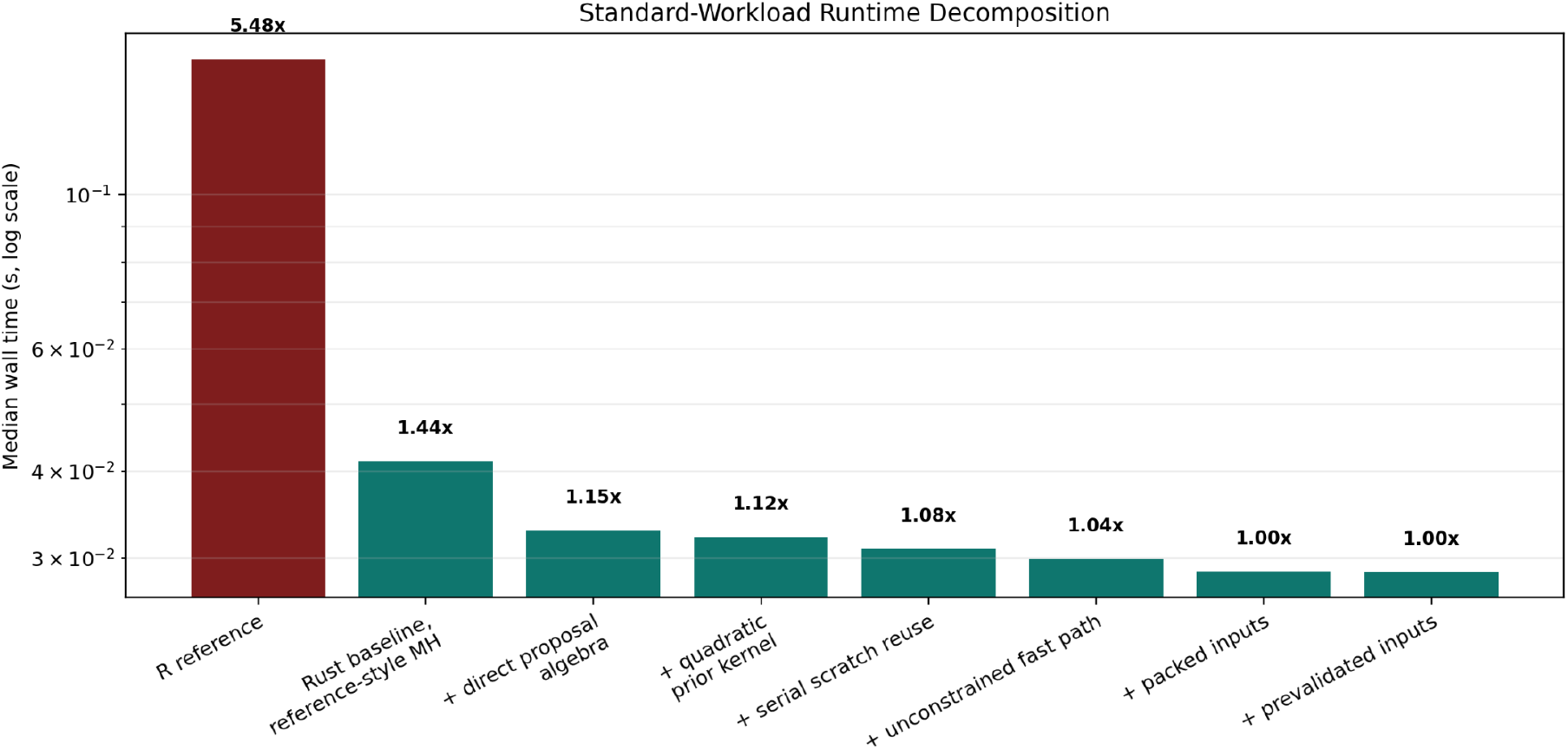
Bars represent sequential implementation stages, with wall-time multiples indicated relative to the previous stage. The R reference is the original bayesm package.

Our primary test of the standard rewrite was the digital-camera conjoint replication^5^. The dataset contained 332 respondents, 16 choice tasks per respondent, and five alternatives per task; we fit it with rhierMnl-RwMixture. Table G1 and Figure G2 report the timing differences. All implementations were timed on the same machine. At *R* = 100,000 Markov chain Monte Carlo (MCMC) draws, the rewrite was 2.31*×* faster single-threaded and 4.35*×* faster on eight threads; at *R* = 200,000, it was 2.71*×* and 9.51*×* faster. All experiments also met the stated population-mean agreement criterion: the largest scaled population-mean gaps were 0.177 single-threaded and 0.043 on eight threads at *R* = 100,000, and 0.237 and 0.049 at *R* = 200,000. We used long MCMC chains so that startup costs did not drive the comparison.

To further evaluate the correctness and performance of our rewrite, we reproduced the results from the marketing study by Dubé et al.,^6^ which uses bayesm extensively to fit a demand-estimation model. We found that the numerical results met the study’s stated agreement criteria and observed a 2–3*×* wall-clock speedup when bayesm-rs was used. Similarly, we evaluated the digital-camera hierarchical logit mixture benchmark described in *Bayesian Statistics and Marketing*, using longer chains so that sampler startup costs did not drive the comparison. bayesm-rs reproduced the same posterior summaries with a 2.7*×* speedup in the single-threaded setting and a 9.51*×* speedup with eight threads.

**Table G1.**
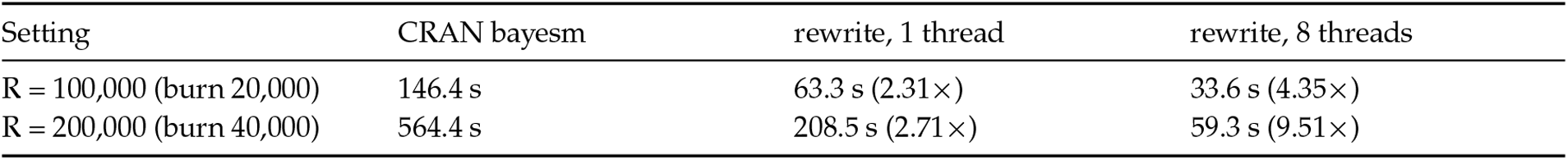
Camera timing against the standard CRAN bayesm 3.1.7 (keep = 5). Parentheses indicate the speedup over the reference.

| Setting | CRAN bayesm | rewrite, 1 thread | rewrite, 8 threads |
| --- | --- | --- | --- |
| $R = 100,000$ (burn 20,000) | 146.4 s | 63.3 s (2.31×) | 33.6 s (4.35×) |
| $R = 200,000$ (burn 40,000) | 564.4 s | 208.5 s (2.71×) | 59.3 s (9.51×) |

**Figure G2.**
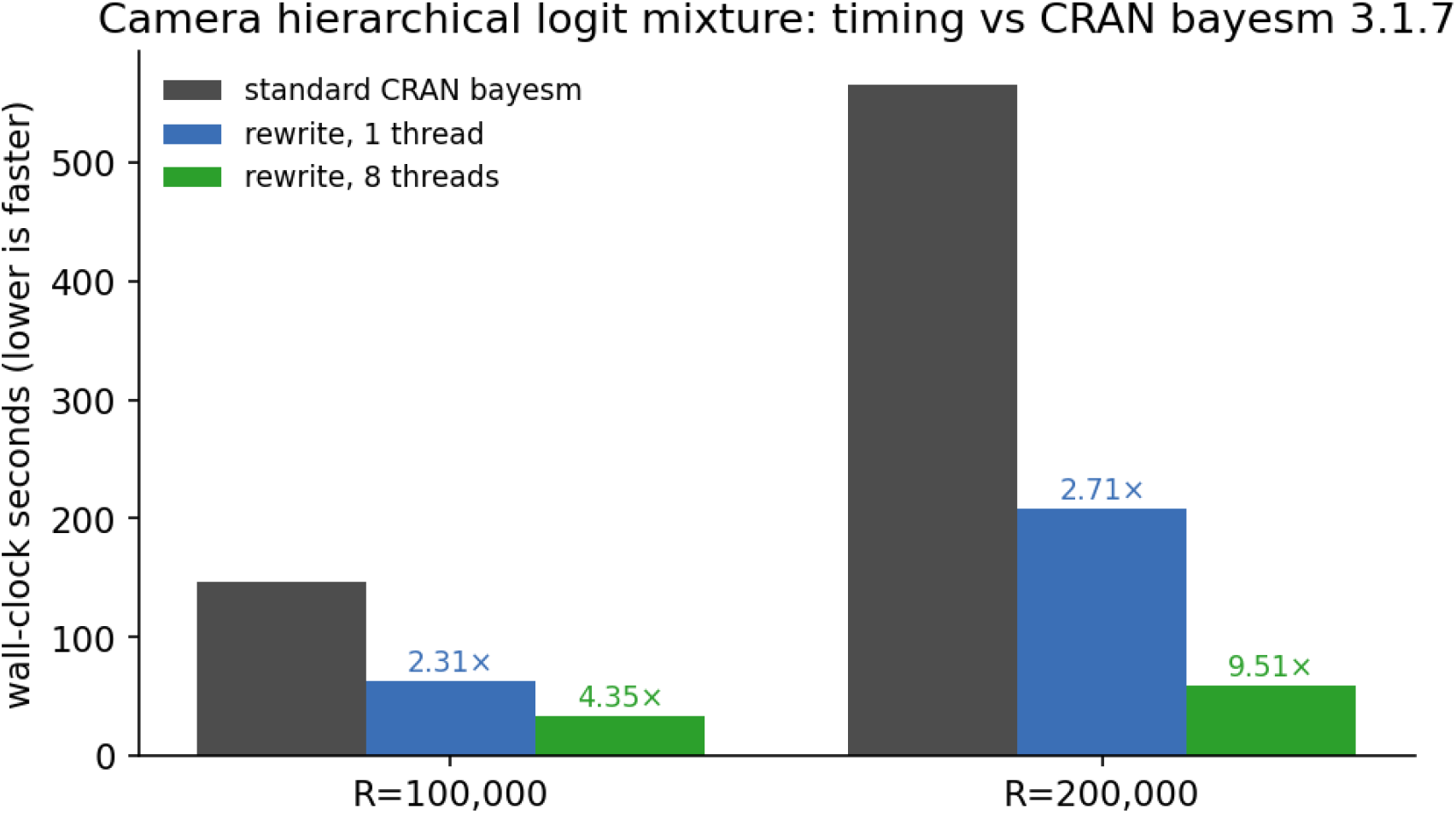
Camera replication timing against CRAN bayesm 3.1.7 (lower is faster).

Our most extensive testing focused on rhierMnlRwMixture, but we also evaluated several other functions. Table G2 reports single-threaded timings from runs with chain lengths chosen for each function so that startup costs were negligible. All runs met the population-mean agreement criterion.

**Table G2.** Single-threaded timing for further functions, same protocol as Table G1.

| Function (dataset) | R | CRAN bayesm | rewrite | speedup |
| --- | --- | --- | --- | --- |
| `rhierNegbinRw` (detailing) | 10,000 | 102.1 s | 27.4 s | 3.73× |
| `rhierLinearModel` (cheese) | 100,000 | 8.3 s | 2.0 s | 4.11× |
| `rbprobitGibbs` (simulated) | 30,000 | 13.8 s | 5.6 s | 2.48× |
| `rivGibbs` (simulated) | 50,000 | 29.6 s | 5.1 s | 5.83× |

#### Initial HMC/NUTS validation

The results in this section are from the first-pass gradient engines; the corrected-validation section reports the corresponding results after correction.

We first tested the new samplers on a simulated hierarchical MNL with four coefficients, using the standard random-walk sampler as the reference and comparing both gradient modes, HMC and NUTS, against it. We used simulated data so that we could control the problem’s size and dimensionality. All three samplers met the convergence requirements. The gradient-based samplers met the population-mean agreement criterion, and NUTS produced 1.37*×* the reference sampler’s effective samples per second. Table G3 presents the exact figures.

**Table G3.**
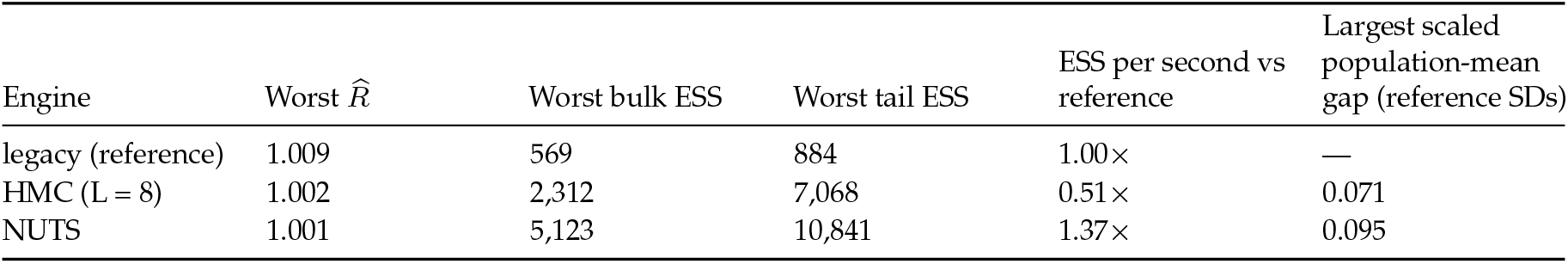
Four-coefficient hierarchical MNL, *R* = 8,000 post-warmup draws after 2,000 warmup iterations. For fixed-trajectory HMC, *L* denotes the number of gradient steps per trajectory. Chains were required to meet the convergence criteria before comparison: rank-normalized split *R̂* below 1.01 and bulk and tail ESS above 400 for every parameter (*R̂* compares between- and within-chain variance; values near 1 indicate agreement among chains).

| Engine | Worst $\hat{R}$ | Worst bulk ESS | Worst tail ESS | ESS per second vs reference | Largest scaled population-mean gap (reference SDs) |
| --- | --- | --- | --- | --- | --- |
| legacy (reference) | 1.009 | 569 | 884 | 1.00× | — |
| HMC ($L = 8$) | 1.002 | 2,312 | 7,068 | 0.51× | 0.071 |
| NUTS | 1.001 | 5,123 | 10,841 | 1.37× | 0.095 |

To test scaling, we then doubled the number of coefficients to eight, using 100 respondents and 20 tasks per respondent. Random-walk mixing degraded as dimensionality increased. Both gradient modes outperformed the reference sampler in ESS per second: HMC achieved 1.83*×* and NUTS achieved 1.45*×*, while both met the population-mean agreement criterion (Table G4, Figure G3).

**Table G4.**
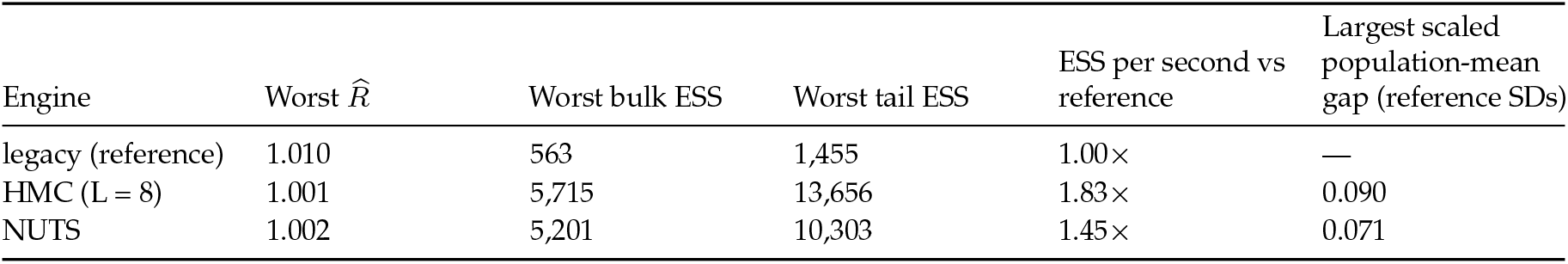
Eight-coefficient hierarchical MNL, *R* = 8,000 post-warmup draws. The convergence requirements were the same as in Table G3.

**Figure G3.**
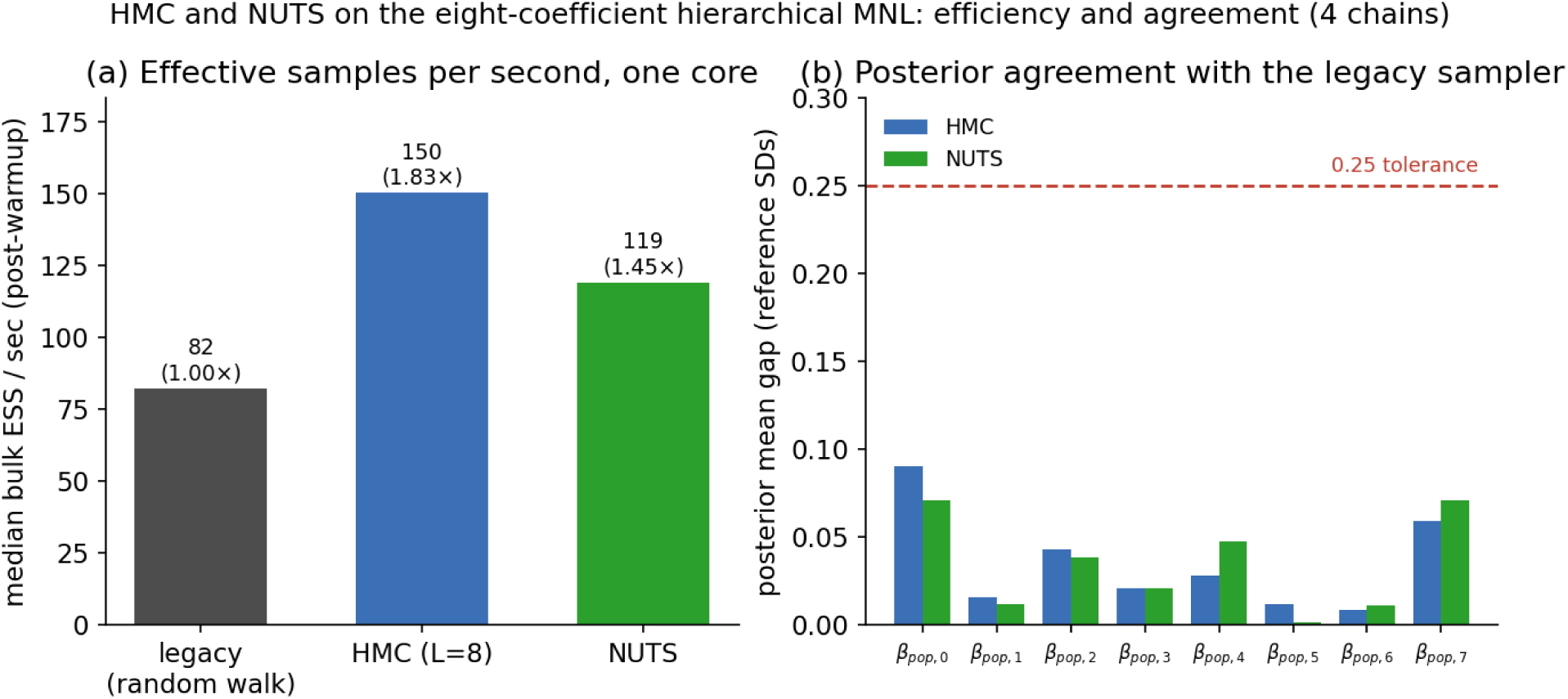
Eight-coefficient hierarchical MNL comparison of the reference, HMC, and NUTS samplers. (a) Median bulk ESS per second after warmup. (b) Absolute scaled gap of each population mean relative to the converged reference sampler.

We ran the same comparison on a second model, the individual negative-binomial model (rnegbinRw), again using simulated data and the same convergence requirements. Neither gradient mode converged. The reference sampler converged at *R* = 16,000, but NUTS yielded a bulk ESS of 224 and a split *R̂* of 1.018 for log_alpha, while HMC failed the *R̂* criterion for two beta coordinates and reported bulk ESS values near 3.6 times the number of draws. These mixed results suggested that the problem was not the likelihood code itself but the adaptation machinery used by the gradient samplers. We later traced the failure to the mass-matrix adaptation.

The HMC/NUTS option was also implemented for three additional functions: rmnlIndepMetrop, rhierNeg-binRw, and rhierMnlDP. All three implementations met the initial population-mean agreement criterion, but testing indicated that gradient sampling offered no advantage for these functions, which matched the theoretical expectation.

#### Initial bayesm.HART validation

Before correction, we compared the HART rewrite with the bank-data example from bayesm.HART’s own documentation (946 respondents, 14 coefficients, *R* = 5,000), using the same data, covariates, and prior settings. The rewrite’s respondent-by-coefficient prediction surface had a correlation of 0.991 with that of the original bayesm.HART implementation and captured most of the focal segment separation (1.91, compared with 2.10 for the original and 0.11 for a linear model). However, 11 of 14 coefficients had cross-respondent mean gaps above 0.25 reference SDs, with the focal Out_State coefficient having a signed difference of *−*1.20. In addition, the rewrite ran 3.6*×* slower than the original sampler: 50.0 s versus 13.9 s single-threaded, and the eight-thread run was no faster. Thus, the first-pass rewrite preserved much of the qualitative segmentation behavior but was not a faithful drop-in replacement for bayesm.HART. The next section investigates the causes, and the corrected-validation section reports the corrected results.

### Obstacles and failure modes

In contrast, some additional human guidance was needed to achieve the current level of performance. Interestingly, GPT-5.2 was not very good at overall prioritization; for example, it would spend a large amount of effort testing increasingly marginal optimizations for one function, without realizing that very low-hanging fruit existed for other bayesm functions. In this case, manual prompting was necessary to ‘redirect’ the model’s attention to other functions. Another example of the model’s deficiencies in higher-level planning comes from its benchmarking efforts. At some point in the rewrite process, the model was asked to write R and Python bindings for the underlying Rust code; however, it ran incomparable, separate sets of benchmarks to compare the performance of the Rust functions with the R or Python bindings. Again, manual prompting was required to directly instruct the model to unify its benchmarking of the different code paths.

Overall, the bayesm-rs rewrite project proceeded mostly autonomously, although there were several critical junctures at which it appeared that GPT-5.2 had gotten ‘stuck’ pursuing some rather myopic approaches (e.g. failing to divert optimization efforts to other functions). Fewer than five hours of actual engineer time were required to prompt or guide GPT-5.2, which is quite impressive given the complexity of the source package. We suspect that newer models, such as GPT-5.6 Sol, would require significantly less intervention, as newer GPT models appear much stronger at working autonomously for hundreds of hours without losing track of higher-level goals.

Neither extension was correct on the first pass. In the HMC/NUTS implementation, the diagonal mass matrix was inverted, and simulation-based calibration later exposed a trajectory-construction defect. The first-pass bayesm.HART rewrite ran 3.6*×* slower than the original because one routine rebuilt a large matrix every iteration, and it applied a single fixed shrinkage constant where the original adapts shrinkage to each coefficient’s scale. The base bayesm rewrite was more successful, but these failures show that AI-assisted package rewriting still requires careful statistical validation. The preceding sections describe the initial results; the following sections diagnose the defects and report the corrected implementations.

As evidenced by the extensions, package rewriting with AI agents is not foolproof. Both extensions contained defects in their first-pass implementations. The following subsections diagnose these defects and draw lessons for practitioners rewriting statistical packages.

#### Mass matrix inversion

The mass-matrix defect was relatively straightforward. HMC needs an estimate of how wide each parameter’s posterior is so that it can take large steps on wide parameters and small steps on narrow ones. The function that supplied that estimate, OnlineVar::inv_mass(), returned the opposite of what was needed. In this implementation, the quantity named inv_mass is used internally as the coordinate-wise scale for position updates, so the required value is proportional to posterior variance rather than precision. The correct implementation returns approximately the posterior variance, but the function was erroneously written to return one divided by the variance.

This defect caused the rnegbin failure. With the scales reversed, the sampler treated the widest parameter as the narrowest and the narrowest as the widest. In rnegbin, the widest posterior belonged to log_alpha, which was about nine times wider than the beta coefficients, so it received the smallest steps and barely moved over the whole run. The beta coefficients correspondingly received steps that were too large and bounced from one side of the posterior to the other on consecutive draws, an alternating pattern that caused the ESS estimator to report values several times the number of draws.

Two checks confirmed this. First, the width estimate learned for each parameter was one divided by that parameter’s true posterior variance, the opposite of what was needed (Figure G4). Second, instructing the sampler to skip the learned widths allowed it to take steps 30 times larger. It then came close to converging with only 4,000 draws (Table G5).

**Table G5.** Individual negative-binomial model (rnegbinRw), using the same simulated data as the initial validation; 4,000 post-warmup draws per run.

| Run | adapted step size | Worst $\hat{R}$ | Worst bulk ESS |
| --- | --- | --- | --- |
| legacy reference (R = 24,000) | — | 1.005 | 482 |
| NUTS, diagonal metric | $4.8 \times 10^{-4}$ | 1.071 | 24 |
| HMC, diagonal metric (L = 8) | $4.3 \times 10^{-4}$ | 1.031 | 92 |
| NUTS, unit metric | $1.4 \times 10^{-2}$ | 1.013 | 368 |

The hierarchical MNL runs in the initial HMC/NUTS validation passed with the same bug active because their parameters all have roughly the same posterior width. When every width estimate is wrong by the same factor, the step-size adaptation absorbs the error.

#### Trajectory construction

Simulation-based calibration (SBC) revealed a second defect in the NUTS implementation, this time in its trajectory construction. The population-mean comparisons in the initial HMC/NUTS validation did not detect the resulting bias because the affected quantity was a hierarchical MNL covariance parameter. The tilted rank histogram in Figure G5 shows the bias before correction; after the trajectory calculation was fixed, the ranks were consistent with uniformity. This result illustrates why agreement on selected posterior means alone was not sufficient to establish sampler correctness.

#### HART runtime slowdown

The HART rewrite’s slowdown arose from an existing routine in bayesm, draw_delta, which updates the coefficients linking respondent characteristics to preferences and then rebuilds a large matrix from scratch at every iteration. When integrating bayesm.HART with bayesm, the agents reused this routine. Its computational cost grows with the square of the number of columns in the respondent-characteristic design matrix *Z*. The original bayesm uses only a few columns, so this cost was negligible. In our comparison,

**Figure G4.**
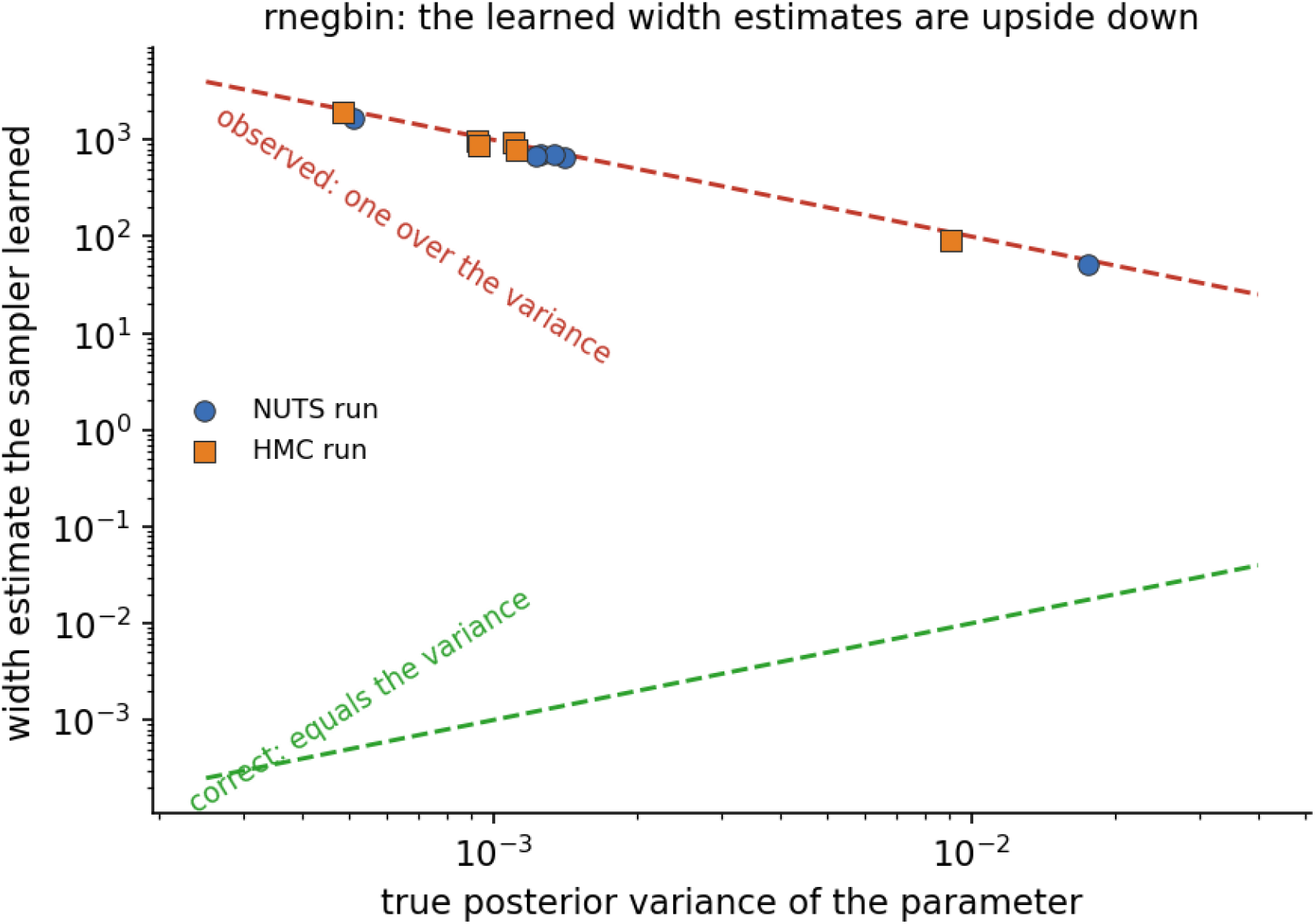
Adapted inverse metric against the posterior variance of each coordinate, log-log, for both diagonal-metric runs: every point falls on the 1/var line (inverted) instead of the var line (correct).

**Figure G5.**
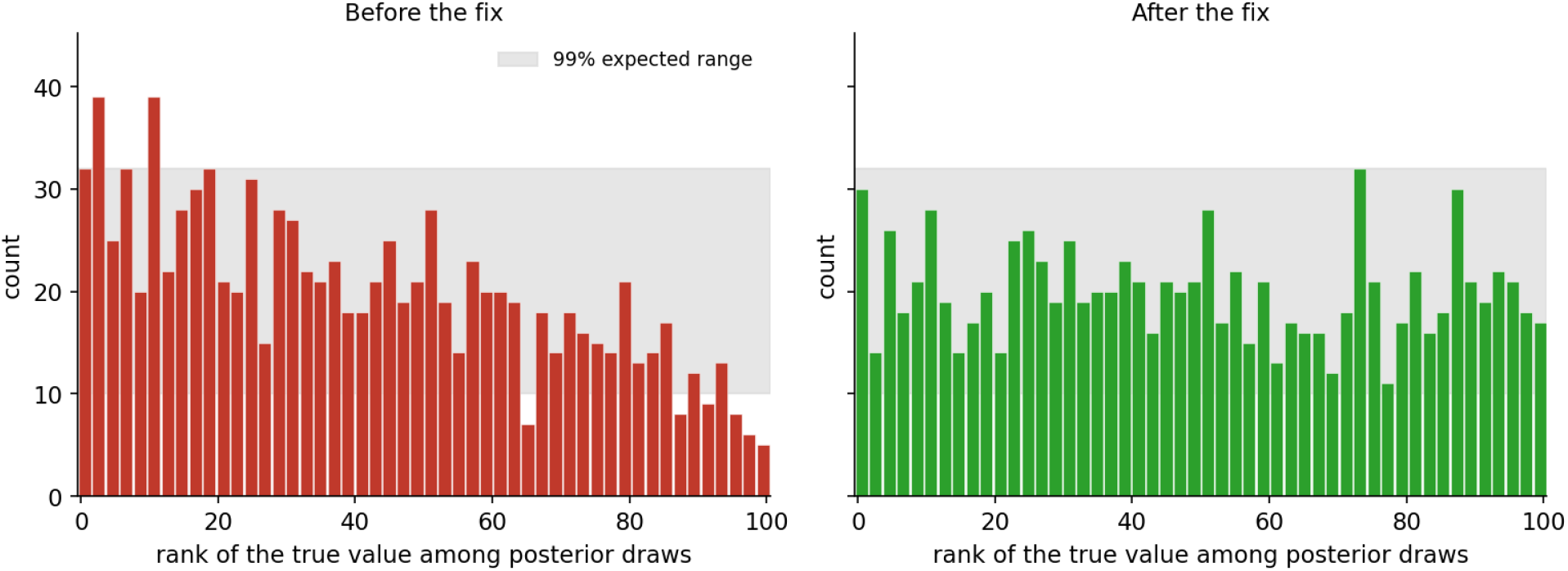
Simulation-based calibration ranks for a hierarchical MNL covariance parameter under NUTS before (left) and after (right) correction of the trajectory-construction defect. The tilted pre-correction histogram indicates systematic bias; the approximately flat post-correction histogram is consistent with uniform ranks.

HART expanded *Z* to 44 columns, sharply increasing the cost. Timing the same model while changing only the column count isolated the issue. At 44 columns, the fit ran about 15*×* slower than the corresponding linear version (Table G6), accounting for the runtime deficit relative to the original HART implementation.

**Table G6.** HART runtime as the number of basis columns grows. The 30-tree row is the configuration used in the initial HART comparison.

| Configuration | basis columns | runtime |
| --- | --- | --- |
| linear Z (5 columns) | — | 0.31 s |
| HART, 5 trees | 6 | 0.33 s |
| HART, 15 trees | 25 | 1.70 s |
| HART, 30 trees (the initial HART configuration) | 44 | 4.71 s |
| HART, 60 trees | 94 | 21.1 s |

#### Prior scaling issues

The second HART defect concerned how strongly the tree effects were shrunk toward zero. In the original HART, the shrinkage adapts: the prior on the tree effects is tied to the covariance matrix that the model is currently estimating, so each coefficient is shrunk in proportion to its own scale. The first-pass rewrite instead used one fixed shrinkage constant, leaf_precision, for every coefficient, so there was no adaptation.

Rerunning the bank comparison with that constant set tighter or looser makes this easy to see (Figure G6). The amount of heterogeneity the model explains moves accordingly: *R*^2^ of 0.063, 0.152, and 0.272 for tighter, default, and looser, bracketing the original HART’s 0.214. The shrinkage gap in the initial HART validation is a result of this constant being wrong, and no single fixed value can be right for every coefficient. The agents made a design error here: where the original lets the estimated covariance matrix set each coefficient’s shrinkage automatically, the rewrite collapsed that rule into one hand-set number. Adding more trees could not fix this: a 60-tree version explains less (*R*^2^ 0.129) and takes 9.4*×* longer.

**Figure G6.**
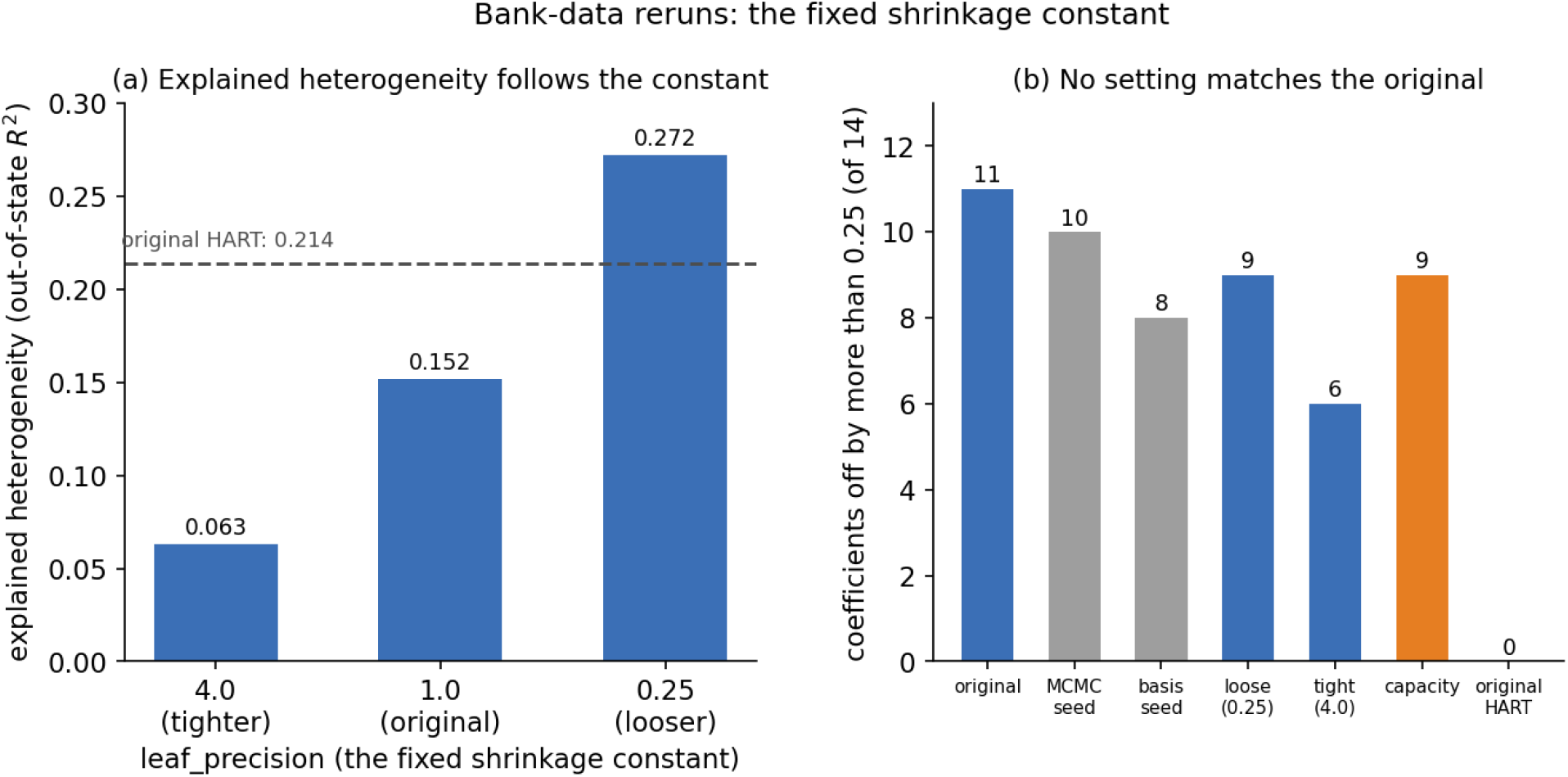
The bank-data reruns. (a) Explained heterogeneity (out-of-state *R*^2^) as the fixed shrinkage constant varies, with the original HART’s value shown by the dashed line. (b) Number of coefficients with gaps exceeding the 0.25-SD tolerance in each rerun: changing seeds alone changes the count (gray), no setting matches the original count of zero, and the 60-tree version (orange) does not help.

### Corrected validation and evidence

After making these corrections, we validated the rewritten samplers using the diagnostics described by Vehtari et al.^4^ Table G7 summarizes the before- and after-correction results.

**Table G7.**
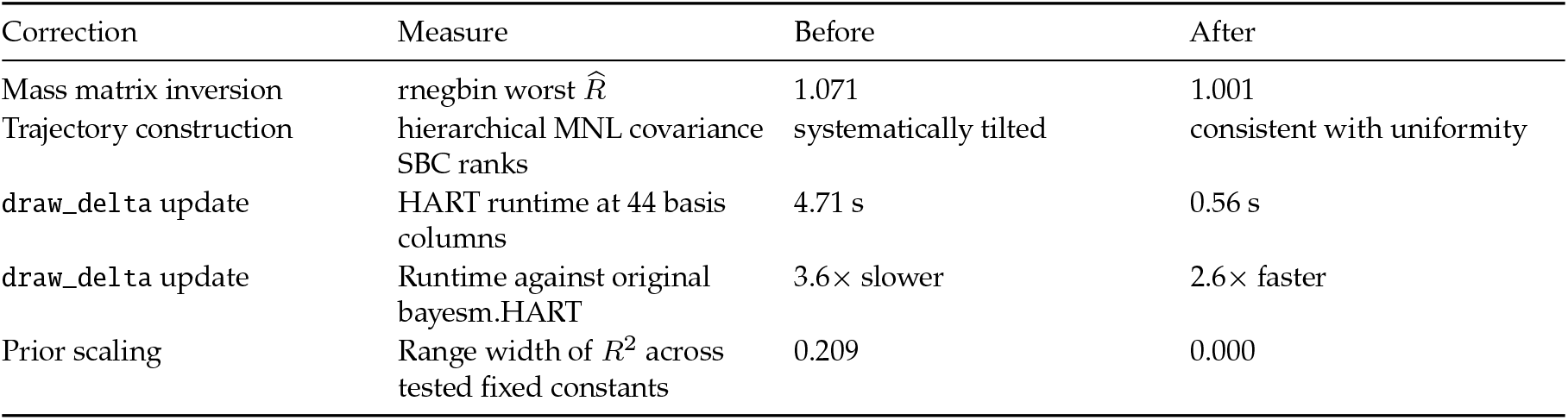
Summary of the principal first-pass defects and their corrected results.

After fixing the mass matrix, the negative-binomial model converged. The gradient-based samplers met the population-mean agreement criterion relative to the reference sampler (largest scaled gap 0.10), and NUTS produced 1.89*×* the reference sampler’s effective samples per second. Fixed-trajectory HMC produced draws that alternated around the posterior mean, inflating the standard effective-sample estimate; after capping ESS at the number of draws, HMC achieved 5.2*×* the reference sampler’s ESS per second. We also reran the hierarchical MNL comparisons from the initial HMC/NUTS validation using the corrected engines. At eight coefficients, HMC achieved 1.60*×* and NUTS achieved 1.72*×* the reference sampler’s ESS per second.

**Table G8.**
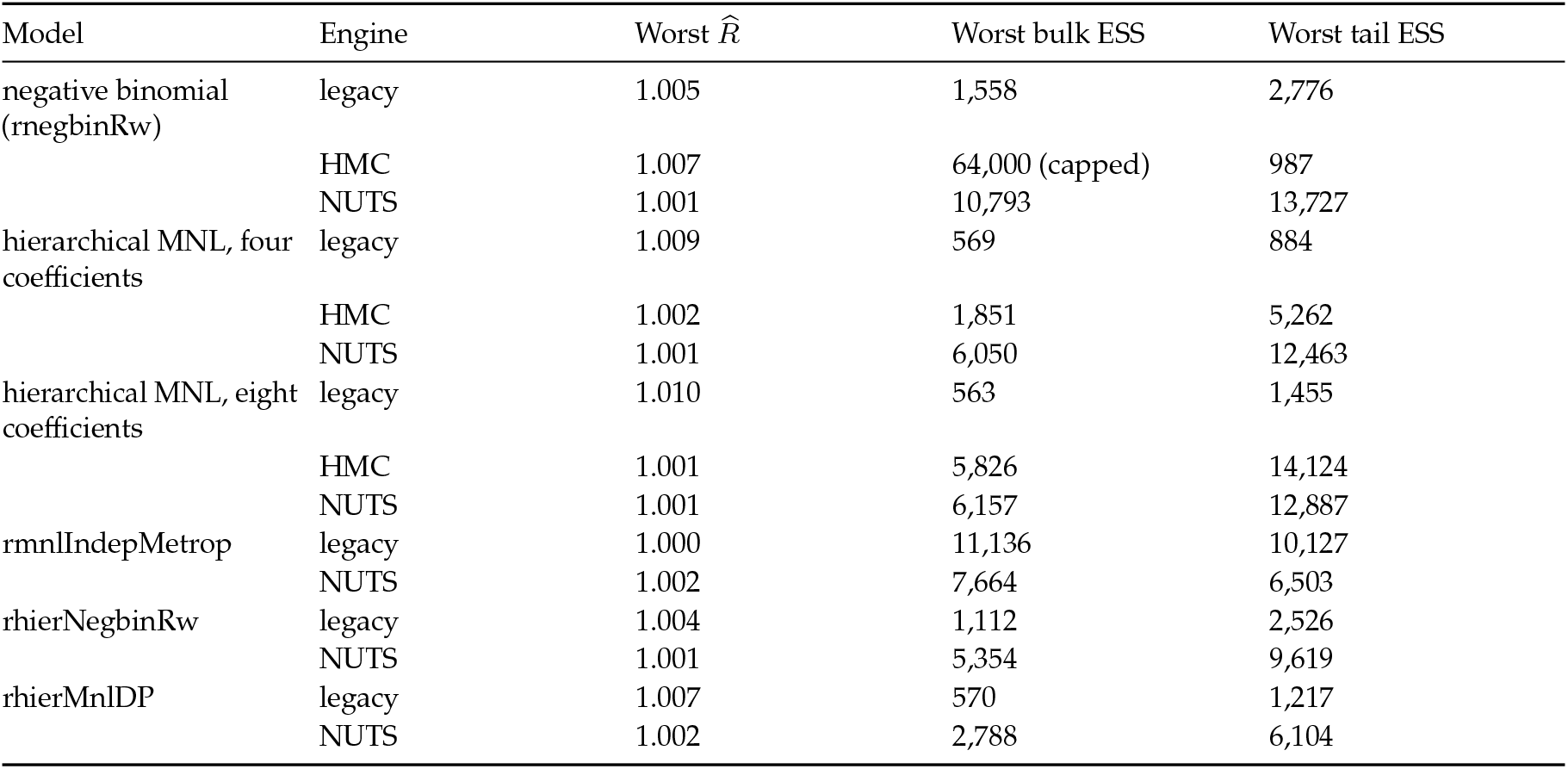
Convergence diagnostics for every model evaluated with the corrected engines: four chains per run, rank-normalized split *R̂*, and bulk and tail effective sample sizes, following Vehtari et al.^4^ Every run met the requirements (*R̂* below 1.01 and both ESS measures above 400). HMC’s alternating draws inflated the standard bulk ESS estimate for the negative-binomial model, so the reported bulk ESS was capped at the number of draws.

The draw_delta update now precomputes the portions that do not change across iterations. At the 44-column configuration, the fit takes 0.56 s rather than 4.71 s. The corrected rewrite is now about 2.6*×* faster than the original bayesm.HART implementation. The HART prior now restores coefficient-specific scaling through the current covariance matrix Σ. After that correction, changing the obsolete global leaf_precision constant no longer changes the out-of-state *R*^2^.

**Figure G7.**
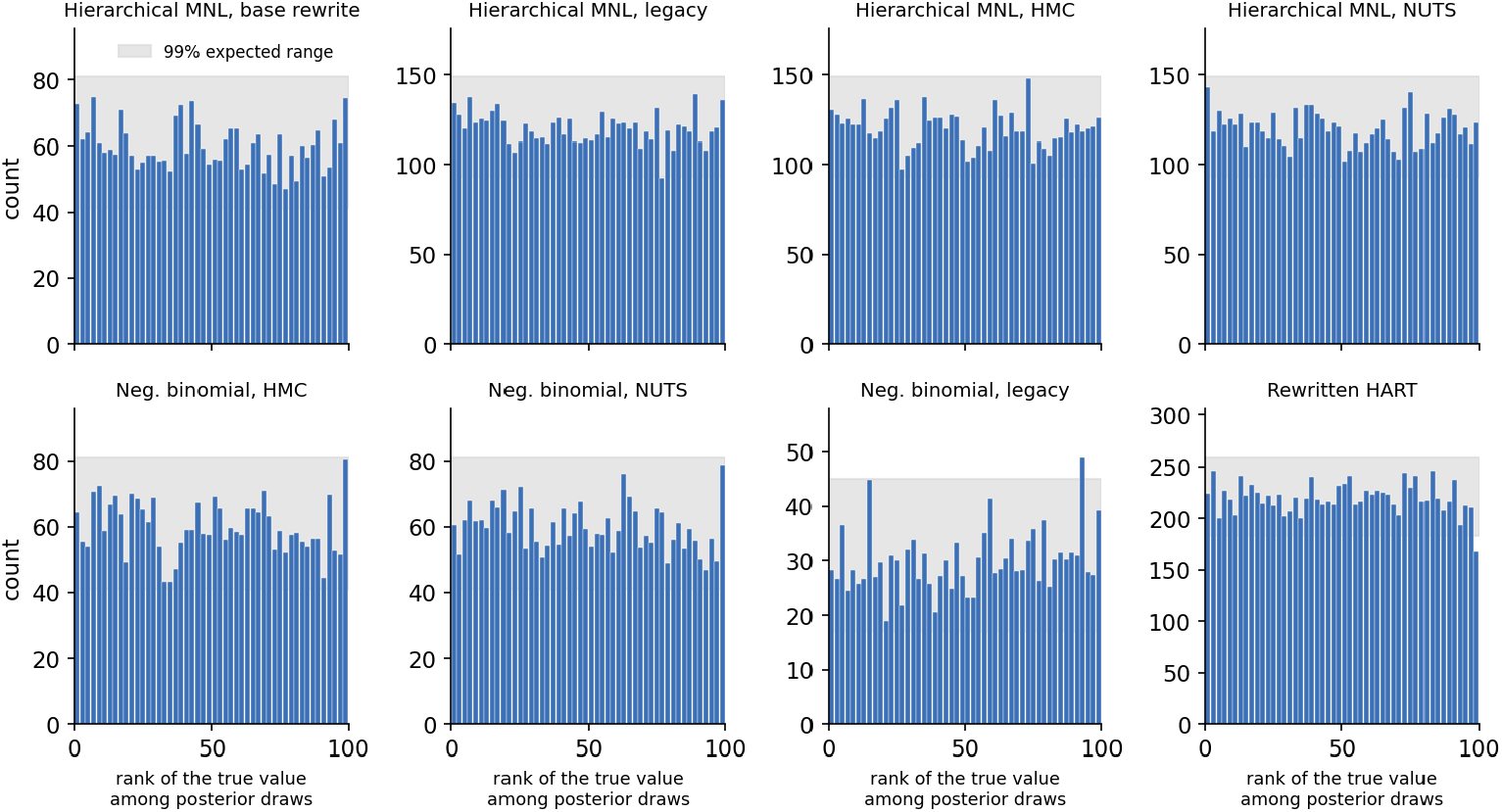
Rank histograms for the validated samplers in this study, pooling each sampler’s checked parameters. The gray band is the range expected from uniform ranks; a correct sampler stays within it.

As a final check, we validated every corrected sampler in this study using the SBC procedure of Talts et al.^7^. The procedure repeatedly draws a true parameter value from the model’s prior, simulates data from it, runs the sampler on the simulated data, and records where the true value ranks among the posterior draws. If the sampler is correct, these ranks are uniform, and the shape of any deviation indicates the kind of error. Across 500 to 1,000 replications per model, the rank histograms for the corrected samplers were consistent with uniformity (Figure G7).

### Outcome

We rewrote bayesm in Rust and showed that bayesm-rs met the population-mean agreement criterion in the digital-camera replication while running faster in both single- and multithreaded settings. We then added HMC/NUTS and HART extensions and subjected them to population-mean agreement checks, convergence diagnostics, and simulation-based calibration. Neither extension was correct on the first pass. The HMC/NUTS engine contained an inverted mass matrix and a trajectory-construction defect, while the HART rewrite contained a quadratic-time draw_delta update and omitted the coefficient-specific Σ scaling used by the original method. The first-pass HART output nevertheless had a 0.991 correlation with the original prediction surface, illustrating how apparently strong aggregate agreement can coexist with important implementation defects.

After correction, all tested samplers met the stated convergence requirements and produced rank histograms consistent with uniformity. On the eight-coefficient hierarchical MNL, fixed-trajectory HMC and NUTS achieved 1.60*×* and 1.72*×* the reference sampler’s effective samples per second. On the negative-binomial model, NUTS achieved 1.89*×* the reference sampler’s effective samples per second. The corrected HART rewrite was 2.6*×* faster.

### Lessons and reflections

The relatively straightforward structure of the bayesm library—independent functions that can be compared with their original implementations—means that post-release maintenance is expected to be limited.

That being said, the ability of GPT-5.2 to completely port a statistical library from one language to another—in this case including fairly complex mathematical operations—illustrates well the strengths of modern coding agents.

The agent mistakes followed a clear pattern. The base rewrite had a more complete reference implementation: each function could be compared directly with CRAN bayesm. However, reference outputs did not fully specify the coding decisions required for the extensions. Practitioners using coding agents for statistical package rewrites should assess both whether the code can be checked against an objective reference and whether it depends on unstated implementation choices.

### Artifacts

|  |  |
| --- | --- |
| Rewritten code | [ajh-oai/bayesm](https://github.com/ajh-oai/bayesm) |
| HART extension | HART extension draft pull request #1: https://github.com/ajh-oai/bayesm/pull/1 |
| HMC/NUTS | HMC/NUTS extension draft pull request #2: https://github.com/ajh-oai/bayesm/pull/2 |

### References

1. Wiemann, T. bayesm.HART. https://github.com/thomaswiemann/bayesm.HART.

2. Hoffman, M. D., & Gelman, A. (2014). The No-U-Turn Sampler: Adaptively setting path lengths in Hamiltonian Monte Carlo. *Journal of Machine Learning Research*, 15, 1593–1623.

3. Rossi, P. E., Allenby, G. M., & McCulloch, R. (2005). *Bayesian Statistics and Marketing*. Wiley.

4. Vehtari, A., Gelman, A., Simpson, D., Carpenter, B., & Bürkner, P.-C. (2021). Rank-normalization, folding, and localization: An improved R-hat for assessing convergence of MCMC (with discussion). *Bayesian Analysis*, 16(2), 667–718.

5. Allenby, G. M., Brazell, J., Howell, J., & Rossi, P. E. (2014). Economic valuation of product features. *Quantitative Marketing and Economics*.

6. Dubé, J.-P., Fang, Z., Fong, N., & Luo, X. (2017). Competitive Price Targeting with Smartphone Coupons. *Marketing Science*, 36(6), 944–975.

7. Talts, S., Betancourt, M., Simpson, D., Vehtari, A., & Gelman, A. (2020). Validating Bayesian inference algorithms with simulation-based calibration. arXiv:1804.06788.

### CASE STUDY H

### Agentic optimization of a shotgun read simulation library

**Andrew Ho^1^ Gene Myers^2^**

^1^ OpenAI; ^2^Diploid Genomics, Inc.

### Summary

GPT-5.2, when prompted in a zero-shot manner to identify potential optimizations to the HI.SIM genomic read simulation library, was able to identify a series of additive, low-level changes resulting in a 23.72% runtime reduction on a reference workload while preserving byte-level output equivalence. A follow-up optimization pass using GPT-5.6 identified an additional series of optimizations resulting in a further 9.5% runtime reduction.

### Scientific context

HI.SIM is an existing suite of tools written in C that allows a user to generate a model of error types and read lengths for a dataset of shotgun reads (HImodel) and simulate reads with matching error and read length distributions from a source genome (HIsim).

### Problem or opportunity

As part of a collaboration between the J. Craig Venter Institute and OpenAI, we explored the feasibility of agentically driven optimization of the read simulation algorithm.

### Agentic intervention

The changes made to the HI.SIM code were primarily local in nature; rather than making fundamental algorithmic changes to the read simulator or modifying the statistical error-type modeling, they generally reduced or eliminated constant-factor overhead in frequently executed operations. The optimizations made include:

- Adding a fast path for reads without sequencing errors
- Removing repeated floating-point divisions from the mutation loop
- Calculating read-length changes while generating edits, eliminating an extra pass
- Replacing base-by-base sequence copying with memcpy
- Buffering FASTA headers, sequences, and error traces before writing
- Replacing repeated fprintf calls with fewer fwrite operations
- Using lightweight integer-to-string formatting
- Reusing dynamically sized buffers across reads
- Adding large userspace file buffers
- Decoding four packed genome bases per loop iteration
- Reusing the source genome for zero-mutation haplotypes instead of copying it
- Centralizing allocation, file-opening, and error handling
- Fixing an incorrect buffer-growth calculation exposed by insertion-heavy reads

Collectively, these changes can be grouped into three broad categories:

- Elimination of repeated work in inner loops
- Elimination of unnecessary memory movement or allocation
- Batching together large numbers of small writes

Each optimization made was separately benchmarked and verified to produce superior performance relative to the previous baseline.

### Human role

There was essentially no human intervention aside from framing the problem in the initial prompt. The model was instructed to search for, implement, and benchmark optimizations without any constraints specified on the nature of such optimizations.

### Validation and evidence

Altogether, these optimizations result in an aggregate 30.97% runtime reduction on a reference four-workload suite:

- 50 kbp synthetic genome at 1,000*×* coverage
- 2 Mbp synthetic genome at 120*×* coverage
- *E. coli* K-12 MG1655 reference at 80*×* coverage
- *S. cerevisiae* S288C reference at 50*×* coverage

All benchmarking used a synthetic HiFi-like model with a 15 kbp mean read length and 1% mean error rate, and the headline result compares the sum of per-workload median runtimes.

### Outcome

Overall, this work provides a concrete demonstration that coding agents can autonomously identify, implement, benchmark, and validate meaningful performance improvements in specialized scientific software. Two successive zero-shot optimization passes reduced aggregate runtime by 30.97% across a representative four-workload suite while preserving byte-level output equivalence and the simulator’s underlying statistical behavior. These results suggest that systematic agentic optimization could offer a low-cost and readily deployable way to recover substantial cumulative efficiency gains across the long tail of scientific software.

### Obstacles and failure modes

This style of optimization is the most conservative, in that it maximally preserves existing structure and compatibility, as opposed to larger refactorings that change the underlying algorithmic reasoning or port code between languages. It is conceivable that more aggressive changes would result in greater speedups; at the same time, however, conservative changes are the most easily integrated into existing code, especially if the proposed changes do not originate from the original author of the code.

### Lessons and reflections

The successful zero-shot optimization of complex, low-level code in the HI.SIM library while maintaining byte-level output equivalence demonstrates that large language models can be successfully used for performance optimizations in specialized scientific subdomains. In this particular case, we aimed to preserve the basic algorithmic structure of the code, focusing only on smaller, “local” optimizations in the areas of the code which dominated runtime.

The ease with which we were able to attain a *>* 30% runtime reduction on low-level code suggests that a relatively large amount of low-hanging fruit exists in scientific libraries. At a minimum, it appears worthwhile to run zero-shot optimization passes over scientific or numerical libraries prior to publication to identify straightforward improvements. Indeed, identifying and implementing local optimizations draws upon some of the greatest strengths of AI coding agents, which are indefatigable and will happily spend hours upon hours proposing small optimizations and carefully profiling their impact. Given that real-world work is still often constrained by the availability of compute–with commonplace routines in genomics potentially taking many days or longer to run, depending on the size of the input data–small, accretive improvements can compound into significant cost and time savings.

Another strength of the agentic approach is the breadth of optimization hypotheses that can be evaluated economically. In addition to the changes ultimately adopted, the agent investigated a number of other hypotheses such as different compiler flags, link-time and profile-guided optimization, alternative random-number generation, etc. Specialist labor is quite scarce, and prior to the development of coding agents, surveying the search space in such a broad manner would be entirely impractical. The agent also independently constructed benchmark workloads and implemented automations for runtime and memory measurements and regression checks, again to a level of thoroughness that would simply not have been feasible merely several years ago.

### Artifacts

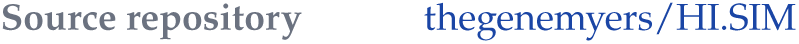

